# Trypanosomatid selenophosphate synthetase structure, function and interaction with selenocysteine lyase

**DOI:** 10.1101/2020.01.28.922716

**Authors:** Marco Túlio Alves da Silva, Ivan Rosa e Silva, Lívia Maria Faim, Natalia Karla Bellini, Murilo Leão Pereira, Ana Laura Lima, Teresa Cristina Leandro de Jesus, Fernanda Cristina Costa, Tatiana Faria Watanabe, Humberto D’Muniz Pereira, Sandro Roberto Valentini, Cleslei Fernando Zanelli, Julio Cesar Borges, Marcio Vinicius Bertacine Dias, Julia Pinheiro Chagas da Cunha, Bidyottam Mittra, Norma W. Andrews, Otavio Henrique Thiemann

**Author notes:** These authors have equal contribution to the paper (Marco Túlio Alves da Silva, Ivan Rosa e Silva).

## Abstract

Early branching eukaryotes have been used as models to study the evolution of cellular molecular processes. Strikingly, human parasite of the Trypanosomatidae family (*T. brucei*, *T. cruzi* and *L. major*) conserve the complex machinery responsible for selenocysteine biosynthesis and incorporation in selenoproteins (SELENOK/SelK, SELENOT/SelT and SELENOTryp/SelTryp), although these proteins do not seem to be essential for parasite viability under laboratory controlled conditions. Selenophosphate synthetase (SEPHS/SPS) plays an indispensable role in selenium metabolism, being responsible for catalyzing the formation of selenophosphate, the biological selenium donor for selenocysteine synthesis. We solved the crystal structure of the *L. major* selenophosphate synthetase and confirmed that its dimeric organization is functionally important throughout the domains of life. We also demonstrated its interaction with selenocysteine lyase (SCLY) and showed that it is not present in other stable complexes involved in the selenocysteine pathway, namely the phosphoseryl-tRNA^Sec^ kinase (PSTK)-Sec-tRNA^Sec^ synthase (SEPSECS) and the tRNA^Sec^-specific elongation factor (eEFSec)-ribosome. Endoplasmic reticulum stress with ditiothreitol (DTT) or tunicamycin upon selenophosphate synthetase ablation in procyclic *T. brucei* cells led to a growth defect. On the other hand, only DTT presented a negative effect in bloodstream *T. brucei* expressing selenophosphate synthetase-RNAi. Although selenoprotein T (SELENOT) was dispensable for both forms of the parasite, SELENOT-RNAi procyclic *T. brucei* cells were sensitive to DTT. Together, our data suggest a role for the *T. brucei* selenophosphate synthetase in regulation of the parasite’s ER stress response.

**Synopsis:** Selenium is both a toxic compound and a micronutrient. As a micronutrient, it participates in the synthesis of specific proteins, selenoproteins, as the amino acid selenocysteine. The synthesis of selenocysteine is present in organisms ranging from bacteria to humans. The protozoa parasites of the Trypanosomatidae family, that cause major tropical diseases, conserve the complex machinery responsible for selenocysteine biosynthesis and incorporation in selenoproteins. However, this pathway has been considered dispensable for the protozoa cells. This has intrigued us, and lead to question that if maintained in the cell it should be under selective pressure and therefore be necessary. Also, since the intermediate products of selenocysteine synthesis are toxic to the cell, it has been proposed that these compounds need to be sequestered from the cytoplasm. Therefore, extensive and dynamic protein-protein interactions must happen to deliver those intermediates along the pathway. In this study we have investigated the molecular and structural interactions of different proteins involved in selenocystein synthesis and describe its involvement in the endoplasmic reticulum protection to oxidative stress. Our results also show how the interaction of different proteins leads to the protection of the cell against the toxic effects of seleium compounds during selenocysteine synthesis.

## Introduction

*Trypanosoma brucei*, *Trypanosoma cruzi* and *Leishmania sp*. protozoan parasites [1] are collectively responsible for thousands of productive life years lost worldwide, as a consequence of human sleeping sickness [2], Chagas’ disease [3], and leishmaniasis [4], respectively. They cycle between an insect vector and a mammalian host, progressing through different life-cycle stages with varying metabolism, cell morphology and surface architecture. Adverse environmental conditions such as nutrient deficiency, hypoxia, oxidative stress, pH and temperature variation occur throughout their life cycle [5]. Trypanosomatids depend on dynamic gene expression to regulate their adaptation to stress, differentiation and proliferation, in response to diverse environmental signals within different hosts [5, 6]. Interestingly, gene expression is controlled post-transcriptionally by spliced leader (SL) trans-splicing, RNA editing and mRNA stability [6].

In their life cycle, these parasites are exposed to reactive oxygen species that are controlled by a unique thiol-redox system based on trypanothione reductase, tryparedoxin and tryparedoxin peroxidase [7,8,9]. In contrast, the main redox regulatory enzymes in mammals are thioredoxin and gluthatione reductases, which contain selenocysteine (Sec) in the active site [10]. Remarkably, only three selenoproteins, namely SELENOK, SELENOT and SELENOTryp, have been reported in trypanosomatids to contain Sec-based putative redox centers, as confirmed by ^75^Se-labeled homologs from *T. brucei* [11], *T. cruzi* [12] and *L. donovani* [13]. Selenocysteine biosynthesis and incorporation into selenoproteins require an intricate molecular machinery that is present, but not ubiquitous, in all domains of life [14]. In eukaryotes [14,15,16] it begins with tRNA^[Ser]Sec^ acylation with L-serine by the seryl-tRNA synthetase (SerRS) followed by its conversion to Sec-tRNA^[Ser]Sec^, sequentially catalyzed by phosphoseryl-tRNA^Sec^ kinase (PSTK) and Sec-tRNA^[Ser]Sec^ synthase (SEPSECS). Selenophosphate synthetase (SEPHS) is a key enzyme in the Sec pathway, being responsible for catalyzing the formation of the active selenium donor for this reaction, selenophosphate, from selenide and ATP. Finally, a tRNA^Sec^-specific elongation factor (eEFSec) directs the Sec-tRNA^[Ser]Sec^ molecule to the ribosome in response to an UGA_Sec_ codon, in the presence of a Sec insertion sequence (SECIS) in the mRNA. However, a detailed protein-protein and protein-RNA interaction network and the mechanism of selenoprotein biosynthesis and its regulation in trypanosomatids remain poorly understood [12, 17].

Furthermore, little is known about selenium metabolism itself in trypanosomatids [17]. It has been reported that selenium in trace amounts is essential for various organisms throughout several domains of life, although at high concentration it is cytoxic [17, 18]. Selenocysteine is specifically decomposed by selenocysteine lyase (SCLY) into alanine and selenide, which is potentially reused by selenophosphate synthetase in several eukaryotes, including trypanosomatids [19,20,21]. Mammals conserve two paralogues of selenophosphate synthetase, namely SEPHS1 and SEPHS2. Mammalian SEPHS2 is itself a selenoprotein known to be essential for selenophosphate formation and consequently selenoprotein biosynthesis. In contrast, the SPS1 isoform (SEPHS1) does not conserve a cysteine or selenocysteine residue in the catalytic site and is likely involved in redox homeostasis regulation [22,23,24]. Strikingly, the cellular availability of hydrogen peroxide is altered by SEPHS1 deficiency in embryonic mammalian cells [25]. Our group previously showed that trypanosomatids conserve only one selenophosphate synthetase, SEPHS2 (SPS2), containing a catalytic cysteine [26]. Its function has been related to selenoprotein synthesis [11].

Not only *T. brucei* SEPHS2 (*Tb*SEPHS2) but also PSTK, SEPSECS and eEFSec (*Tb*PSTK, *Tb*SEPSECS and *Tb*eEFSec, respectively) [11] independent knockdowns impair selenoprotein synthesis in the parasite procyclic form (PCF). Interestingly, *Tb*PSTK and *Tb*SEPSECS double knockout cell lines demonstrated that *T. brucei* PCF does not depend on selenoproteins [11]. This result was extended with the observation that *Tb*SEPSECS is not essential for normal growth of the bloodstream form (BSF) *T. brucei* [27] and for its survival in the mammalian host [28]. In addition, a *Tb*SEPSECS knockout cell line did not show any growth disruption upon hydrogen peroxide-induced oxidative stress in PCF [27]. This result also seems to be valid for other trypanosomatids, since *L. donovani* SEPSECS null mutant promastigote (*Ld*SEPSECS) cell lines show normal growth even upon oxidative stress, and during macrophage infection [13]. On the other hand, *Tb*SEPHS2 RNAi cells are sensitive to oxidative stress induced by hydrogen peroxide [29]. The reason why trypanosomatids maintain such complex machinery for selenoprotein biosynthesis remains unclear.

Moreover, the function of Kinetoplastida selenoproteins has not been elucidated yet. Interestingly, *T. brucei* PCF and BSF are sensitive to nanomolar concentrations of auranofin [12, 17], an inhibitor of mammalian selenoprotein biosynthesis. Nonetheless, auranofin did not show any differential effect on *Tb*SEPSECS knockout lines, when compared to the wild type *T. brucei* PCF [27] or the counterpart *L. donovani* amastigote [13]. SELENOTryp (SelTryp) is a novel selenoprotein exclusive to trypanosomatids that contains a conserved C-terminal redox motif, often found in selenoproteins that carry out redox reactions through the reversible formation of a selenenylsulfide bond [12]. On the other hand, mammalian SELENOK/SelK [30] and SELENOT/SelT [31] homologs were recently shown to be endoplasmic reticulum (ER) residents, where they have a role in regulating Ca^2+^ homeostasis. Regulation of ER redox circuits control homeostasis and survival of cells with intense metabolic activity [30, 31]. Chemical induction of ER stress with DTT and tunicamycin in PCF *T. brucei*, but not BSF [32] result in ER expansion and elevation in the ER chaperone BiP, inducing the unfolded protein response (UPR) [33,34,35]. Prolonged ER stress induces the spliced leader RNA silencing (SLS) pathway [34]. Induction of SLS, either by prolonged ER stress or silencing of the genes associated with the ER membrane that function in ER protein translocation, lead to programmed cell death (PCD). This result is evident by the surface exposure of phosphatidyl serine, DNA laddering, increase in ROS production, increase in cytoplasmic Ca^2+^, and decrease mitochondrial membrane potential [34].

Despite the wealth of information on the selenocysteine machinery in Eukaryotes, selenoprotein biosynthesis and function in early branching Eukaryotes, such as trypanosomatids, remain poorly understood. Moreover, little is known about selenium metabolism itself in these organisms. Here, we present a detailed structural, biochemical and functional analysis of trypanosomatid selenophosphate synthetase (*Tb*SEPHS2). *Tb*SEPHS2 crystal structure we determined demonstrates that a conserved fold is important for its function. We also show that *Tb*SEPHS2 interacts with *T. brucei* selenocysteine lyase (*Tb*SCLY) *in vitro* and that they co-purify from *T. brucei* procyclic cell extracts. We further demonstrate that the *Tb*SEPHS2-SCLY binary complex is not part of other stable complexes in the Sec-pathway of *T. brucei*, namely the *Tb*SEPSECS-tRNA^[Ser]Sec^-PSTK and the *Tb*EFSec-tRNA^[Ser]Sec^-ribosome. We also determined that *Tb* SEPHS2 ablation in procyclic, but not bloodstream *T. brucei* cells leads to growth defect in the presence of the ER stressors DTT or tunicamycin. Similarly, SELENOT knock down in *T. brucei* cell lines led to sensitivity to DTT, although SELENOT was found to be dispensable for both PCF and BSF *T. brucei*. Together, our data shed light into the protein assemblies involved in the selenocysteine pathway in *T. brucei* and suggest a possible role for the *T. brucei* selenophosphate synthetase in regulation of the parasite’s ER stress response.

## Results

### The L. major selenophosphate synthetase crystal structure is highly similar to its orthologs, despite sharing low amino acid sequence identity

*T. brucei* and *L. major* selenophosphate synthetases SEPHS2 isoforms have low sequence identity to the well characterized orthologs from *Homo sapiens*, *Aquifex aeolicus* and *Escherichia coli* (42%, 29% and 28%, respectively) (Figure S1). We described the crystallization of *Tb*SEPHS2 and ΔN(69)-*Lm*SEPHS2 elsewhere [36]. Full length *Tb*SEPHS2 structure determination was not successful due to lack of sufficient experimental data, and full length *Lm*SEPHS2 was recalcitrant to crystallization. Here we present the crystal structure of ΔN-*Lm*SEPHS2 (PDB 5L16) at 1.9 A resolution solved by molecular replacement using human SEPHS1 (PDB 3FD5) as a search model. The structure was refined to R_free_/R_work_ of 0.21/0.17 (detailed refinement statistics are shown in Table 1).

**Table 1:** ΔN-LmSEPHS2 crystal structure refinement statistics.

| **Refinement** | ***ΔN-LmSEPHS2*** |
| --- | --- |
| Refinement program | REFMAC 5.8.0135 |
| Total number of atoms | 2,736 |
| Number of amino acid residues | 323 |
| Number of solvent atoms | 293 |
| Ligand | 1 molecule of sulfate ion |
| Resolution range (Å) (completeness) | 1.882 - 40.845 (96.1%) |
| Reflections used in refinement (in cross validation, random) | 33,411 (5%) |
| R<sub>work</sub>/R<sub>free</sub> | 0.1732/0.2131 |
| Fo, Fc correlation | 0.95 |
| **B-factors (Å<sup>2</sup>)** |  |
| All atoms | 27.3 |
| Protein atoms | 18.0 |
| Ligand atoms | 46.5 |
| Water | 52.5 |
| **R.M.S.D** |  |
| Bond lengths (Å) | 0.006 |
| Bond angles (°) | 1.018 |

ΔN-*Lm*SEPHS2 crystallized as a monomer in the asymmetric unit showing a typical aminoimidazole ribonucleotide synthetase (AIRS)-like fold [37], which consists of two α+β domains labeled N- and C-terminal AIRS (AIRS and AIRS_C, respectively) ranging from amino acid residues 74 to 190 and 204 to 384, respectively. The N-terminal AIRS domain folds into a six-stranded β-sheet flanked by two α-helices and one 3_10_-helix, while the AIRS_C domain also presents a six-stranded β-sheet that is flanked by seven α-helices and one 3_10_-helix (Figures 1A and 1B). Overall, the ΔN-*Lm*SEPHS2 monomer is highly similar to its orthologs, as revealed by the root mean square (R.M.S.D.) deviation of main-chain atomic positions between 0.7 Å and 2.2 Å when ΔN-*Lm*SEPHS2 is compared to *H. sapiens* SEPHS1 [22] (0.7 Å and 0.8 Å for PDBs 3FD5 and 3FD6, respectively), *A. aeolicus* SEPHS [38] (1.0 Å for PDBs 2ZAU, 2ZOD and 2YYE) and *E. coli* SEPHS (SelD) [39] (2.2 Å for PDB 3UO0) monomers, respectively (Figure 1C). The main differences occur in loops that are longer in *Lm*SEPHS. Our crystal structure suggests that the conserved AIRS-like fold of selenophosphate synthetases is necessary for their enzymatic mechanism.

**Figure 1:**
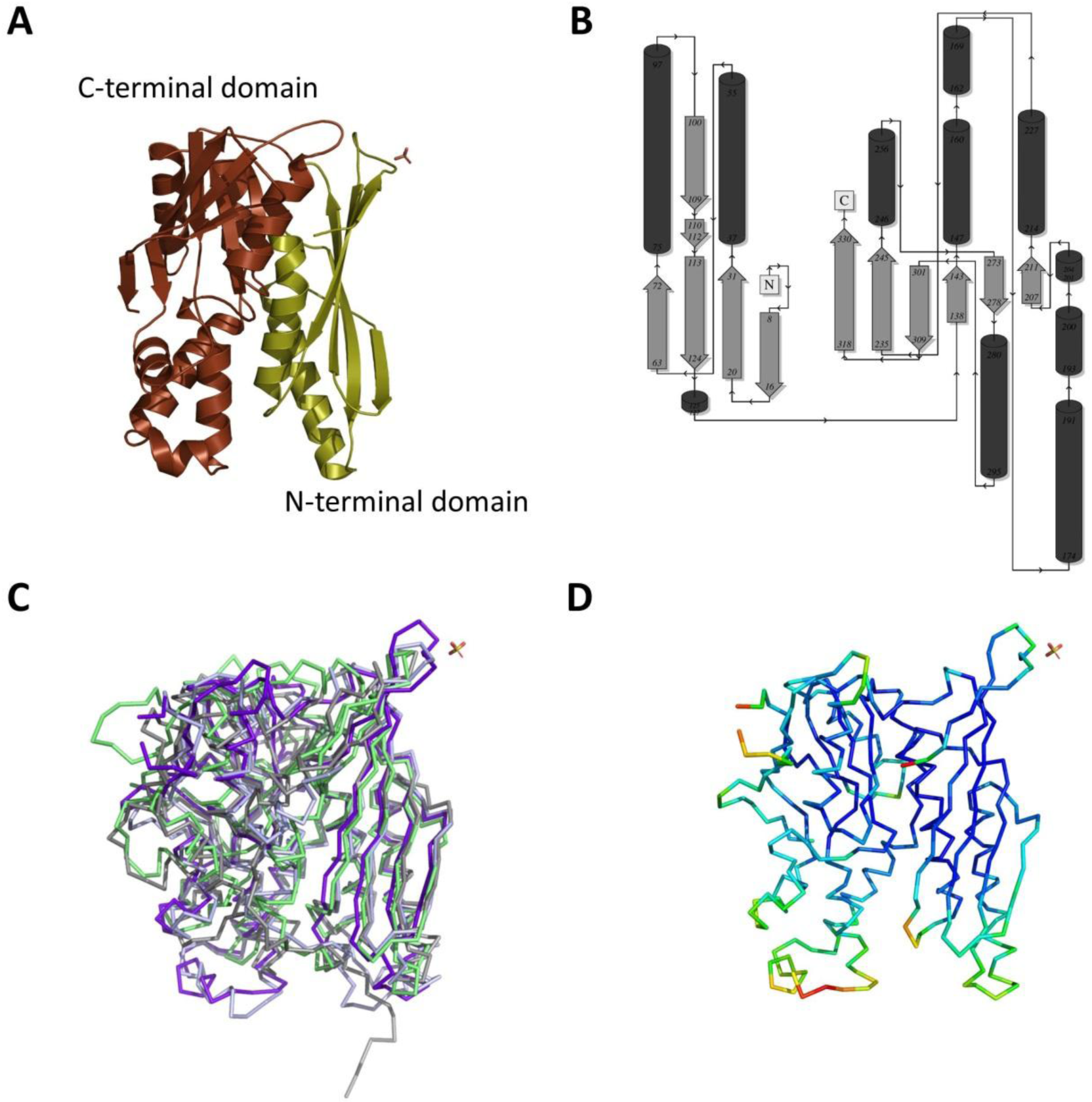
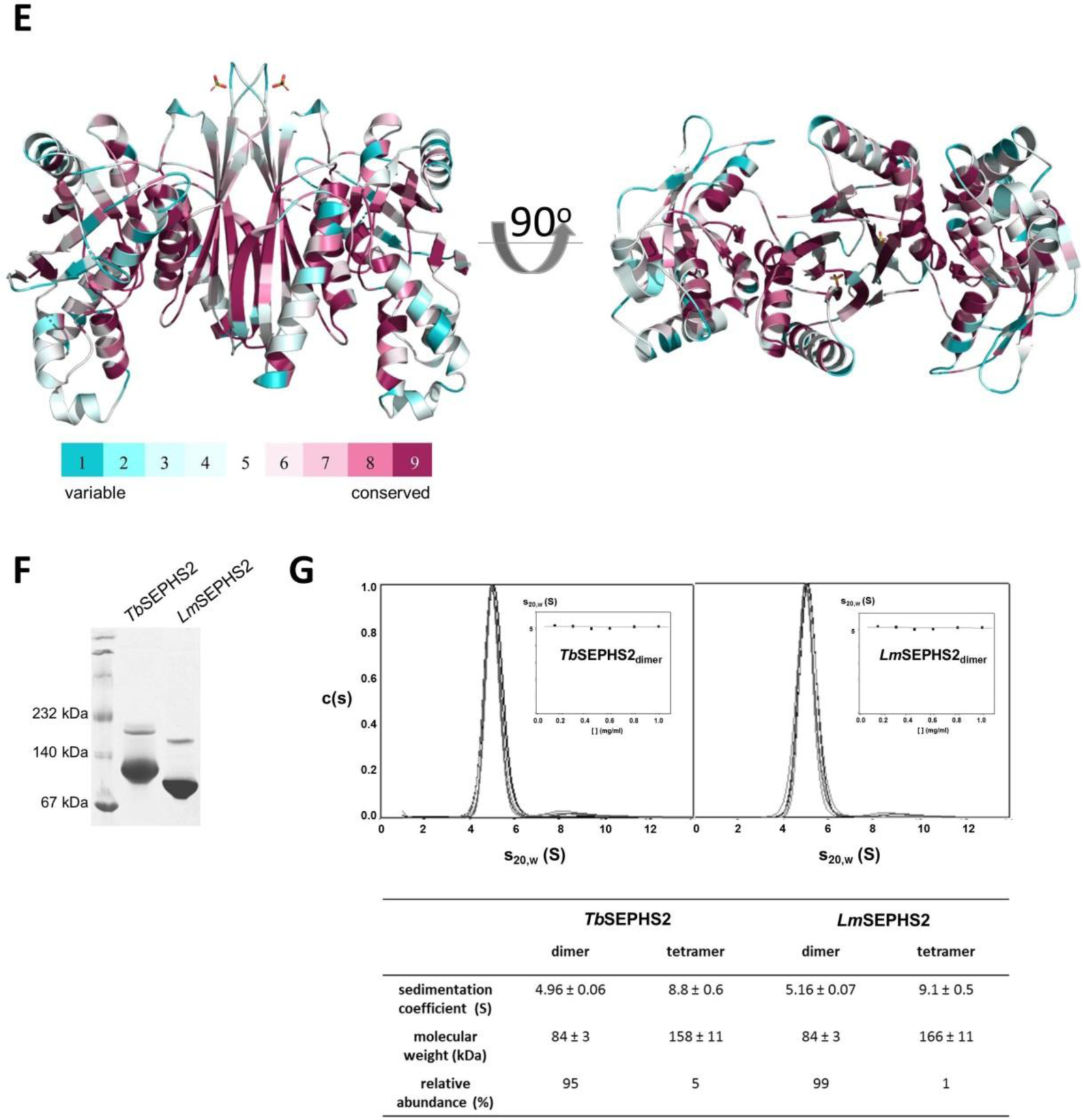
ΔN-*Lm*SEPHS2 crystal structure. **A-** Cartoon representation of the monomer structure in the asymmetric unit showing a typical AIRS-like folding. **B-** Topology factor distribution of the monomer crystal structure. **C-** Superimposition of *Aa*SEPHS (grey) [38], *Ec*SEPHS (green) [39], *Hs*SEPHS1 (light blue) [22] and *Lm*SEPHS2 (purple). **D-** B-factor distribution in the monomer crystal structure (low (blue) to high (red)). **E-** Dimeric model generated using PDBePISA [40] depicting amino acid residue conservation by ConSurf [72]. **F-** Native gel eletrophoresis showing the prevalence of dimers in solution for *T. brucei* and *L. major* selenophosphate synthetases at 2 mg/mL. A small amount of tetramers is also observed for both protein preparations (top bands). **G-** Sedmentation coefficient distribution (c(s)) at increasing protein concentration normalized to the most abundant oligomer (dimer) obtained by sedimentation velocity analytical ultracentrifugation (SV-AUC). SV-AUC data were analyzed in SEDFIT [73]. The insert displays sedimentation coefficients measured for dimers at increasing total protein concentration. The table shows measured sedimentation coeficient, molecular weight and relative abundance of dimers and tetramers.

Notably, selenophosphate synthetases were reportedly active dimers in *E. coli* [39], *A. aeolicus* [38] and *H. sapiens* [22]. Indeed, both recombinant full length *Lm*SEPHS2 and *Tb*SEPHS2 predominantly oligomerize as 83±3 kDa dimers *in vitro* as shown by sedimentation velocity analytical ultracentrifugation (SV-AUC) (Figures 1G and 1H) and native gel electrophoresis (Figure S2). Curiously, a relatively small amount of tetramers was also detected *in vitro* (Figures 1G and 1H). Tetramer-dimer dissociation constants of 161±10 μM and 178±10 μM were measured by sedimentation equilibrium AUC (SE-AUC, Figure S3) for *Tb*SEPHS2 and *Lm*SEPHS2, respectively. The data indicate that the dimer corresponds to the likely dominant form of selenophosphate synthetase in solution. A dimer model of ΔN-*Lm*SEPHS2 (Figure 1E) was generated using PDBePISA [40] as a likely quaternary structure, stable in solution with a 3230 Å^2^ buried surface. The dimerization surface occurs mainly between the β2 and β5 strands of adjacent monomers and is also stabilized by hydrophobic interactions between side chains, leading to the formation of an eight-stranded β-barrel. The dimeric structure of ΔN-*Lm*SEPHS2 conserves two symmetrically arranged ATP-binding sites, formed along the interface between AIRS and AIRS-C domains in each monomer. Amino acid residues previously described to bind ATP phosphate groups [22,38,39] are also conserved (Lys46, Asp64, Thr95, Asp97, Asp120, Glu173 and Asp279), although Lys46 and Asp64 are not present in the crystal structure (Figures 1 and S1).

The N-terminal portion of selenophosphate synthetases has been shown to be highly flexible in the absence of ligand [22,38,39,41,42], and it is disordered in the crystal structure of the apo *A. aeolicus* SEPHS [42]. Similarly, the apo ΔN-*Lm*SEPHS2 crystal structure lacks a 69-amino acid residues-long N-terminal region that includes a glycine-rich loop, where the conserved catalytic residues Cys46 and Lys49 are located. A molecular tunnel formed by the long N-terminal loop in substrate-bound selenophosphate synthetase structures is believed to protect unstable catalysis intermediates [22,38,39]. Interestingly, a novel sulfate binding site was identified in the ΔN-*Lm*SEPHS2 monomer at His84 and Thr85 (Figures 1 and S1).

### The N-terminal region of trypanosomatid SEPHS2 is important but not essential for ATPase activity in vitro and for selenoprotein biosynthesis in selD deficient E. coli

Our group previously showed that both *Tb*SEPHS2 and *Lm*SEPHS2 have a slow kinetics *in vitro* in the presence of selenide [26]. We further evaluated their ATPase activity in the absence of selenide by monitoring the ATP peak over time by HPLC, as shown in Figure 2A. Full-length *Lm*SEPHS2 consumed most of the ATP available *in vitro* during the first five hours of reaction, while full-length *Tb*SEPHS2 consumed half of it during the same period. Interestingly, ΔN(25)-*Tb*SEPHS2, which lacks the predicted disordered N-terminus but preserves all catalytic residues, consumed half of the available ATP only after an 18 hour reaction, indicating that this region is important but not essential for its ATPase activity. On the other hand, ΔN(70)-*Tb*SEPHS2 constructs, which lack the functional residues necessary for selenophosphate formation but conserve most amino acid residues composing the two ATP-binding sites, showed only small residual ATP hydrolysis in the absence of selenide. Curiously, ΔN-*Lm*SEPHS2 showed residual ATPase activity comparable to ΔN(25)-*Tb*SEPHS2. As a control, ATP did not show any residual hydrolysis in the absence of selenophosphate synthetase even after 72 hours of incubation in the reaction buffer.

**Figure 2:**
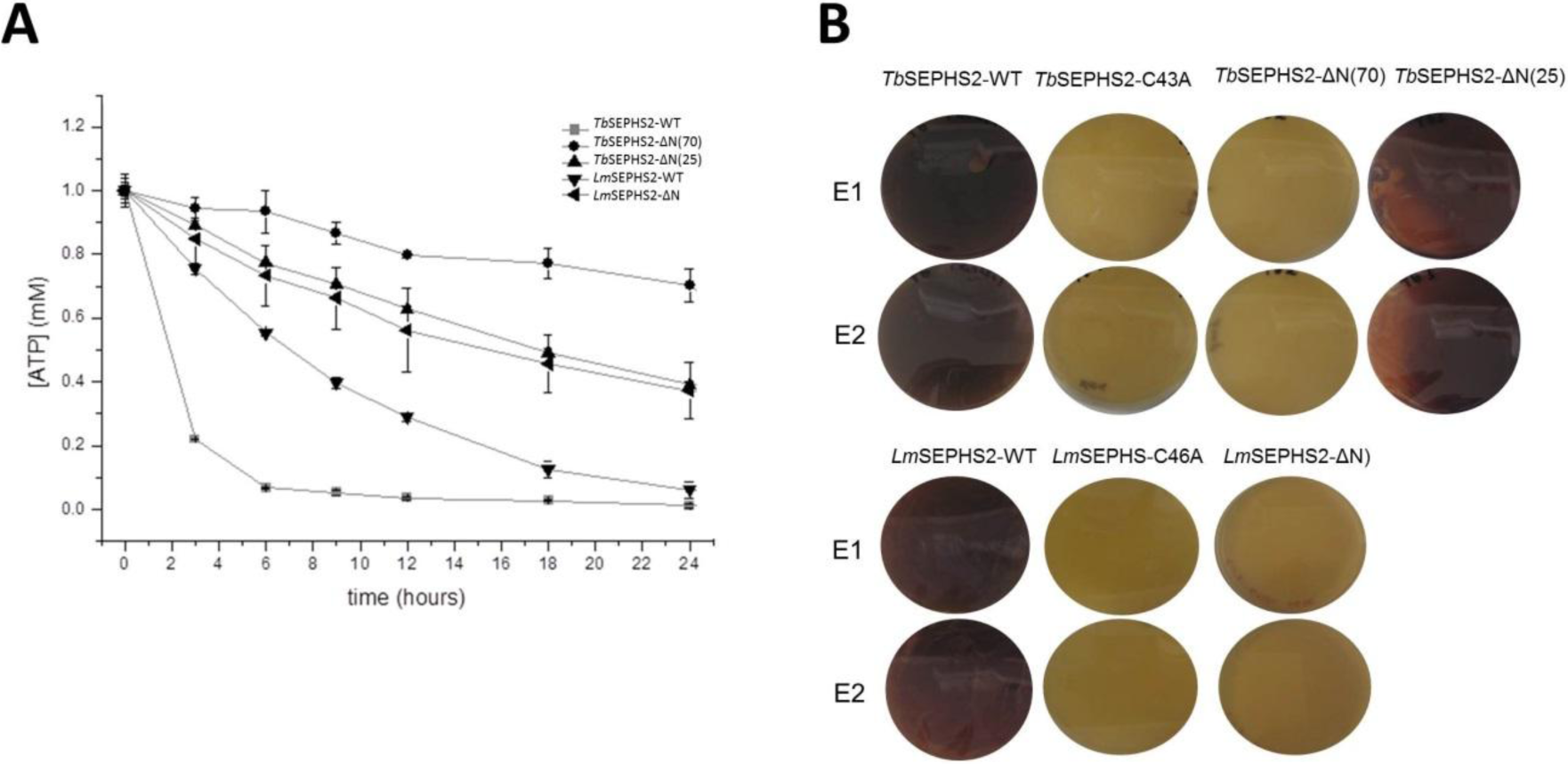
ATPase activity and functional complementation assays. **A-** ATP hydrolysis *in vitro* over time measured by HPLC for full length and N-terminally truncated constructs of *T. brucei* and *L. major* selenophosphate synthetases. **B-** Selenophosphate synthetase functional complementation assays in SEPHS defficient *E. coli* strain (WL400 (DE3)) transformed with different constructs. The purple color indicates a functional formate dehydrogenase H selenoprotein expression. E1 and E2 correspond to biological duplicates.

Like *E. coli* SEPHS (SelD) [39], but in contrast with its *H. sapiens* orthologs, trypanosomatid SEPHS2 is not a selenoprotein itself [26]. *Lm*SEPHS2 was previously shown to restore selenoprotein biosynthesis in a SEPHS deficient *E. coli* WL400(DE3) strain [26]. To extend this information, we verified that both full-length *Tb*SEPHS2 and ΔN(25)-*Tb* SEPHS2, but not ΔN(70)-*Tb*SEPHS2 and ΔN-*Lm*SEPHS2, are also capable of complementing *selD* deletion in *E. coli* (Figure 2B). As expected, no functional complementation resulted from Cys42Ala*-Tb*SEPHS2 and Cys46Ala*-Lm*SEPHS2 mutants, as negative controls (Figure 2B). Notably, although having slow kinetics *in vitro*, ΔN(25)-*Tb*SEPHS2 successfully restored *E. coli* SEPHS function (Figure 2B).

### T. brucei SEPHS2 binds selenocysteine lyase (TbSCLY) but does not co-purify with higher order complexes of the selenocysteine pathway from T. brucei

A putative interaction of eukaryotic selenophosphate synthetase with selenocysteine lyase has been suggested based on reported co-immunoprecipitation of mouse homologs [43]. Thus, we sought to evaluate *T. brucei* SCLY-SEPHS2 direct interaction *in vitro* by sensitive SEC-MALS technique. We unambiguously observed a binary hetero-complex formation *in vitro* (Figure 3A). Isothermal titration calorimetry (ITC) confirmed the interaction (Figure 3B). We also determined that the pyridoxal-phosphate (PLP) molecule bound to the active sites of SCLY is hidden upon SEPHS2 interaction, as measured by a decrease in PLP fluorescence accompanied by a blue shift (Figure 3C). Additionally, we observed that ΔN(70)-*Tb*SEPHS2 does not bind *Tb*SCLY as measured by ITC, indicating that the N-terminal region is necessary for *in vitro* interaction, similarly to the *E. coli* SEPHS (*Ec*SEPHS) N-terminal dependence for SEPHS-SelA-tRNA^[Ser]Sec^ ternary complex formation in *E. coli* [41].

**Figure 3:**
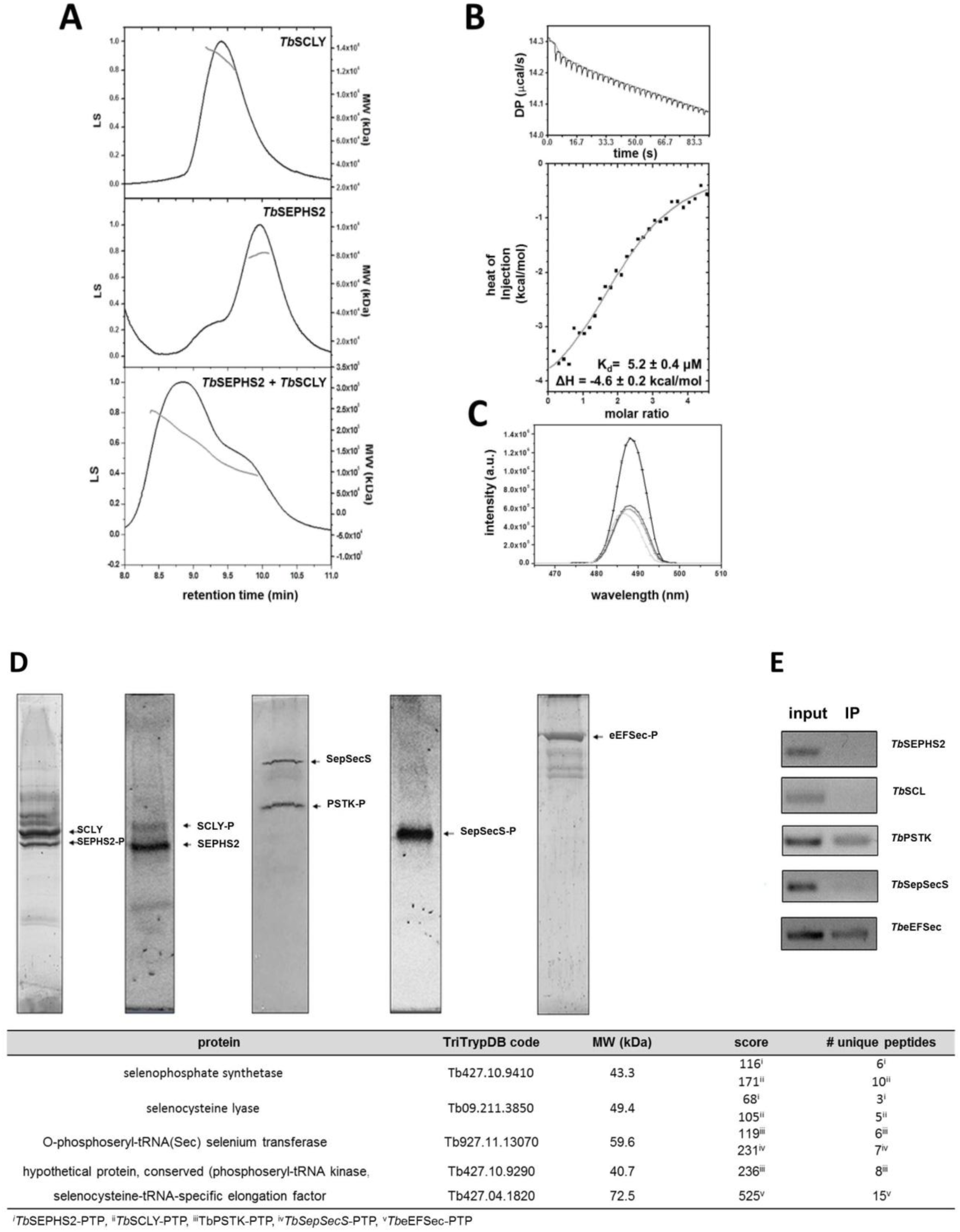
Interaction between selenophosphate synthetase and selenocysteine lyase. **A-** SEC-MALS profiles for *Tb*SCLY, *Tb*SEPHS2 and 1 *Tb*SCLY : 1 *Tb*SEPHS indicating the formation of a binary complex *in vitro*. **B-** ITC curves obtained by *in vitro* titration of *Tb*SEPHS2 to *Tb*SCLY. **C-** PLP-fluorescence in the presence of different concentrations of *Tb*SEPHS2. **D-** SyproRuby^TM^ stained SDS-PAGE of tandem affinity purification (TAP) products of either *Tb*SEPHS2-PTP, *Tb*SCLY-PTP, *Tb*PSTK-PTP, *Tb*SEPSECS-PTP or *Tb*eEFSec-PTP as bait. **E-** Analysis of tRNA^Sec^ copurification by RT-PCR. F-LC-MS/MS analysis of the corresponding SDS-PAGE bands.

Remarkably, *Tb*SEPHS2 and *Tb*SCLY co-purified with each other in independent PTP-TAP experiments (Figure 3D). Also, no tRNA^[Ser]Sec^ was copurified in either experiment, suggesting that this interaction may occur independent of tRNA^[Ser]Sec^. Interestingly, *Tb*SCLY was previously reported to localize predominantly to the nucleus of PCF *T. brucei* [21]. On the other hand, we immunolocalized a C-terminally PTP-tagged construct of *Tb*SEPHS2 both in the nucleus and the cytoplasm of the cell (Figure S6). We further observed that *Tb*PSTK is found both in the cytoplasm and the nucleus of PCF *T. brucei, as Tb*SEPHS2, whereas *Tb*SEPSECS and *Tb*eEFSec are excluded from the nucleus (Figure S6). Since a colocalization experiment was not possible due to the unavailability of antibodies against each protein, we sought to evaluate the formation of putative larger complexes involved in the Sec pathway using PTP (protein A - TEV site - protein C)-TAP (tandem affinity purification) experiments of *Tb*PSTK-PTP, *Tb*SEPSECS-PTP and *Tb*eEFSec-PTP. *Tb*SEPSECS-PTP did not co-purify any other protein, whereas *Tb*PSTK-PTP co-purified with *Tb*SEPSECS (Figure 3D). The C-terminal PTP-tag of *Tb*SEPSECS might have impeded its interaction with *Tb*PSTK. Additionally, no stable protein-protein complex was observed for *Tb*eEFSec-PTP (Figure 3E). RT-PCR analysis revealed that tRNA^[Ser]Sec^ co-precipitates with the *Tb*PSTK-P-SEPSECS complex and with *Tb*eEFSec (Figure 3D). Together, our data demonstrate that no stable complex is formed between *Tb*SEPHS2 and *Tb*PSTK-P-SEPSECS-tRNA^[Ser]Sec^ or *Tb*eEFSec-tRNA^[Ser]Sec^, although these experiments do not exclude the possible formation of transient higher order complexes.

Furthermore, poly-ribosomal profiling experiments showed that neither *Tb*SEPHS2 nor *Tb*SCLY are present in ribosomal complexes involved in the Sec pathway in PCF *T. brucei* (Figure 4A). Conversely, *Tb*PSTK and *Tb*SEPSECS are only present in ribosome-free fractions. A small amount of these proteins was detected in monosome fractions (Figure 4B) possibly due to an overlap between ribosome-free and 40S ribosome fractions, as confirmed by the detection of BiP control in both fractions (Figure 4) consistent with Small-Howard et al [44] results that showed that *Hs*SEPHS1 is also present mammalian in ribosome-free fractions. Additionally, *Tb*eEFSec is mostly present in 80S ribosomes and polysomes, but a small amount of the protein is still found in the ribosome-free fraction and in both 40S and 60S ribosomes (Figure 4C), as shown for *Hs*eEFSec [44]. Disruption of the mounted ribosome with the chelating agent EDTA demonstrated that *Tb*eEFSec dissociates from monosomes or polysomes, being detected only in mRNA-free fractions (Figure 4D).

**Figure 4:**
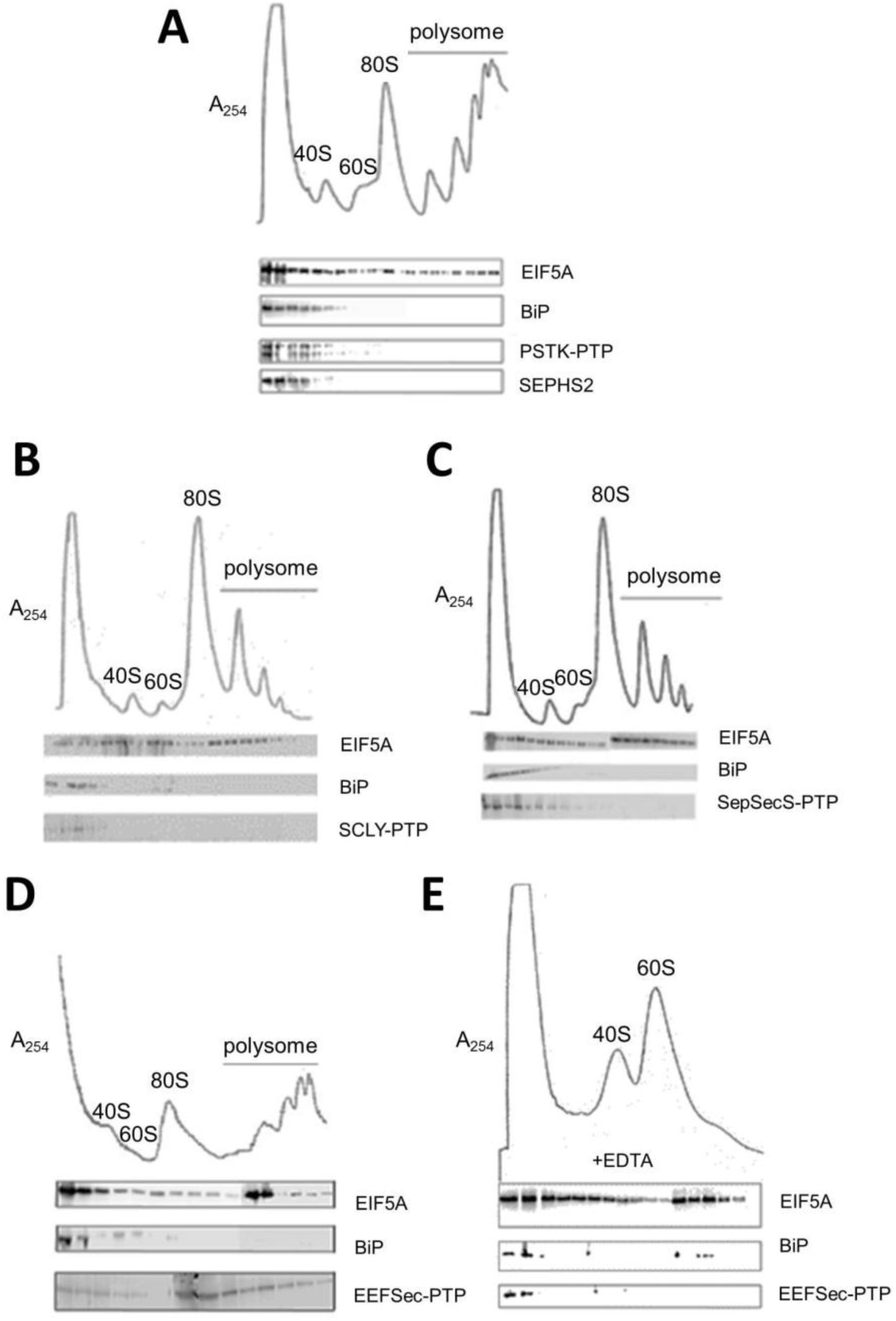
Polysomal profile analysis of selenoproteins synthesis factors. Lysates of PCF *T. brucei* PTP-tagged selenocysteine biosynthesis proteins were fractionated in a sucrose gradient centrifugation (7–47% sucrose) as mRNP (riobosome-free), monosome (40S, 60S and 80S) and polysome fractions as monitored by UV absorbance at 254 nm. Western blot analyses of tagged proteins, using anti-protein A antibody were carried out to localize selenoprotein synthesis factors (**A-** *Tb*PSTK-PTP, **B-** *Tb*SCLY-PTP, **C-** *Tb*SEPSECS-PTP, and **D-** *Tb*eEFSec-PTP). BiP and EIF5A were used as was used as ribosome-free and polysome fraction markers. **E-** Ribosomes dissociation into monosome units in the presence of EDTA fractionated in a 5–25% sucrose gradient.

### *Tb*SEPHS2 RNAi-induced T. brucei cells are sensitive to endoplasmic reticulum chemical stressors

Ablation of selenophosphate synthetase function impairs selenoprotein synthesis not only in mammals [24] but also in *T. brucei* [11]. However, *Tb*SEPHS2 is not essential for either PCF and BSF *T. brucei* under laboratory conditions [11, 27]. Recent work showed that mammalian selenoprotein T (SELENOT) is involved in ER stress response [31]. Therefore, we sought to investigate whether chemical ER stress upon SEPHS2 ablation impairs *T. brucei* viability. We induced RNAi expression against *Tb*SEPHS2 with tetracycline for 48 hours and the cells were subsequently treated with common stressors of ER, namely DTT and tunicamycin (TN) for two hours. TN effectively inhibits the transfer of oligosaccharides onto nascent ER proteins in BSF *T. brucei*, while DTT is thought to generate ER stress by disrupting the redox conditions needed to form protein disulfide bridges in PCF *T. brucei* [33,34,35].

Both DTT and TN caused a slight but significant reduction in viability of induced PCF *T. brucei* cells, suggesting that they negatively interfere with ER metabolism in the absence of SEPHS2 (Figures 5A and 5C). Additionally, SEPHS2-RNAi BSF *T. brucei* cells were induced with tetracycline for 24 hours and subsequently incubated with TN or DTT for two hours. Only the lower concentrations of DTT (150 and 300 µM) tested showed a reduction of SEPHS2-RNAi BSF *T. brucei* cells (Figure 5B) viability. No effect was detected at any concentration of TN in BSF *T. brucei* (Figure 5D). In spite of the lack of transcriptional regulation in *T. brucei*, chemical induction of ER stress apparently results in ER expansion and elevated amounts of the ER chaperone BiP in PCF *T. brucei* [33,34,45]. However, no alteration of BiP expression in both PCF and BSF *T. brucei* cells in the presence of DTT or TN was observed by Western blot (Figures 5I and 5J) despite the negative response to ER stressors. Similarly, Koumandou et al [46] and Tiengwe et al [32] did not observe any change in BiP expression neither at the transcript nor protein level in BSF *T. brucei* upon DTT or TN treatment.

**Figure 5:**
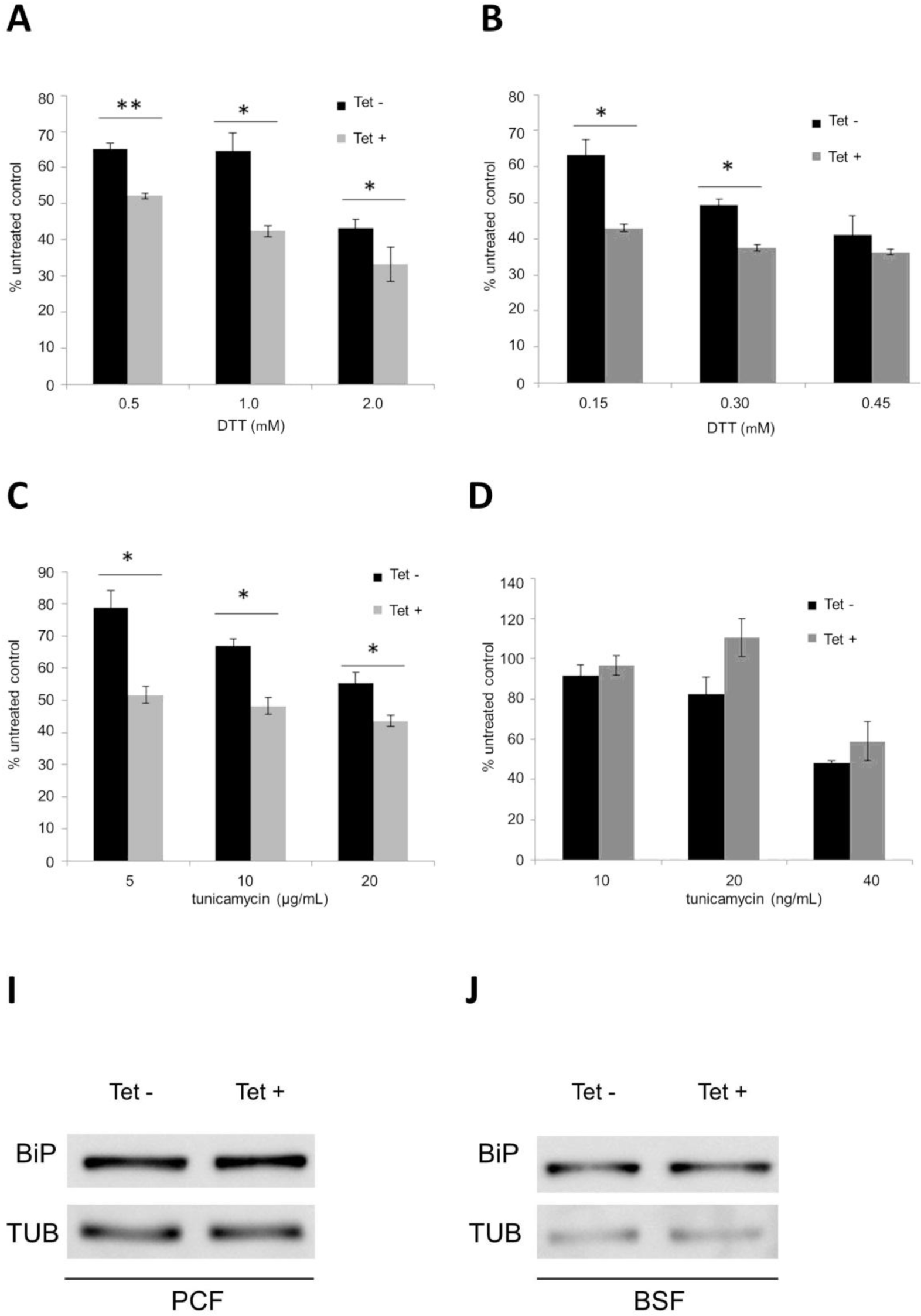
Sensitivity of *Tb*SEPHS2-RNAi *T. brucei* cell lines to DTT or tunicamycin. Tetracycline non-induced (dark bars) and induced (grey bars) *Tb*SEPHS2 RNAi PCF and BSF *T. brucei* cells treated with various concentrations of DTT and tunicamycin. The plots show cell concentration relative to untreated control after a 24h-incubation at 28°C and 37°C for PCF and BSF *T. brucei*, respectively. Bars represent the average of 3 independent experiments including standard deviations of experiments proceeded in **A-** and **C-** PCF *T. brucei* cells, and **B-** and **D-** BSF *T. brucei* cells. The asterisks represent significant differences between the stressors of ER treatment (PCF *T. brucei*: 0.5 mM DTT: ** P = 0.007; 1.0 mM DTT: * P = 0.02; 2.0 mM DTT: P = 0.04; 5μg/mL tunicamycin: * P = 0.033; 10μg/mL tunicamycin: * P =0.01; 20μg/mL tunicamycin: * P = 0.02; BSF *T. brucei*: 0.15 mM DTT: * P = 0.02; 0.3 mM DTT: * P = 0.01; two-tailed Student’s t test). Western blot analysis of BiP in whole cell extracts of **I-** PCF and **J-** BSF *Tb*SPS2 RNAi *T. brucei* cell (12% SDS-PAGE; α-tubulin as a normalization standard).

### Selenoprotein T (SELENOT) is not essential for both procyclic and bloodstream forms of T. brucei, but TbSELENOT RNAi-induced cells are sensitive to endoplasmic reticulum chemical stressors

The mammalian selenoprotein T (SELT, SELENOT) is an ER resident enzyme whose Sec-containing redox domain is believed to regulate various post-translational modifications that require protein disulfide bond formation in the ER including chaperones and also contributing to Ca^2+^ homeostasis [31]. On the other hand, trypanosomatid SELENOT is a selenoprotein whose cellular function has not been characterized yet. Thus, we sought to evaluate whether this enzyme is essential in *T. brucei*. Tetracycline-induced SELENOT-RNAi resulted in 96% reduction of SELENOT mRNA in PCF *T. brucei* as measured by qPCR, but no significant growth defect compared to non-induced cells (Figures 6A and 6C). In BSF *T. brucei*, a slight growth defect was observed for tetracycline-induced cells (Figures 6B and 6D) with around 91% mRNA level reduction.

**Figure 6:**
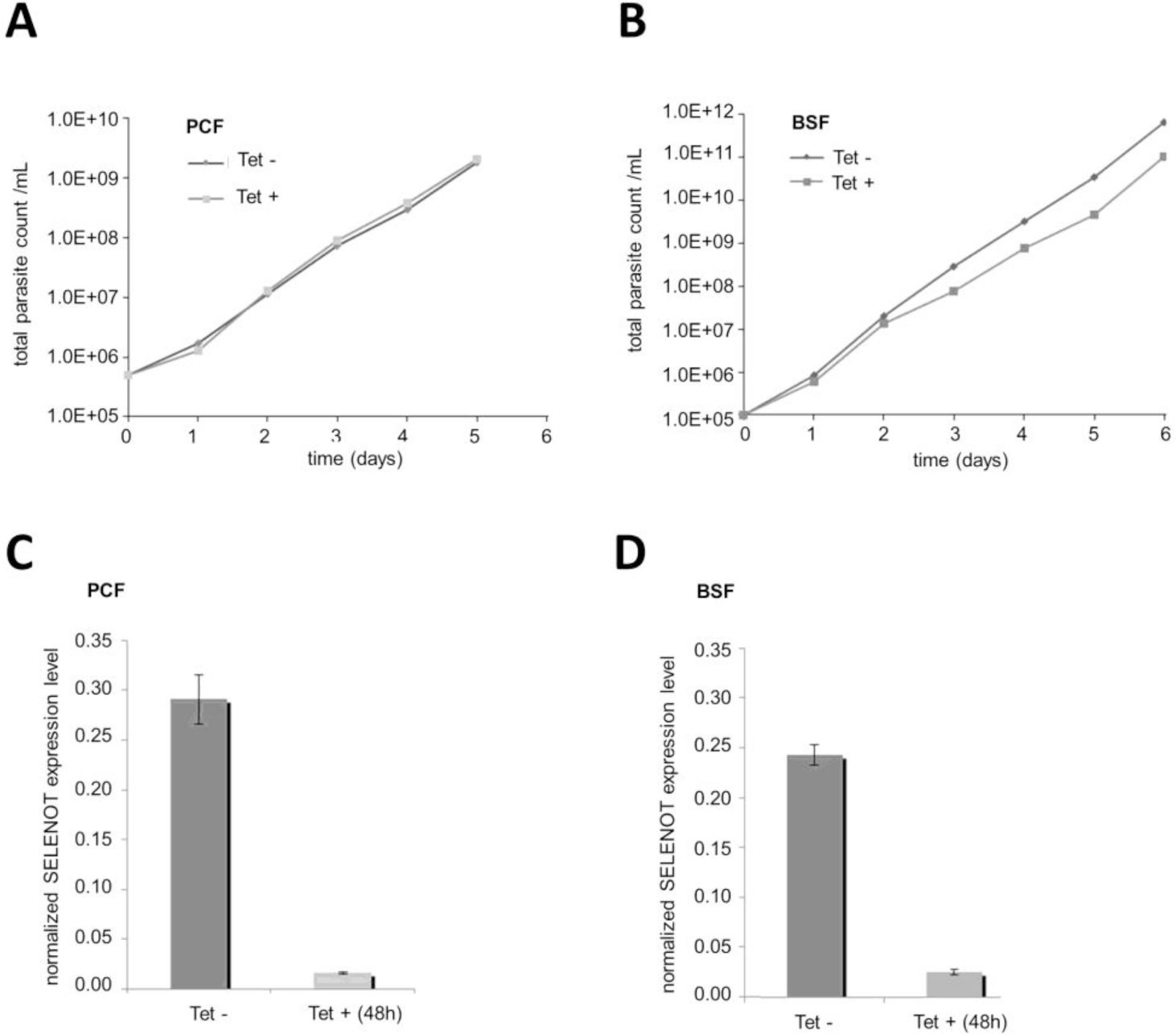
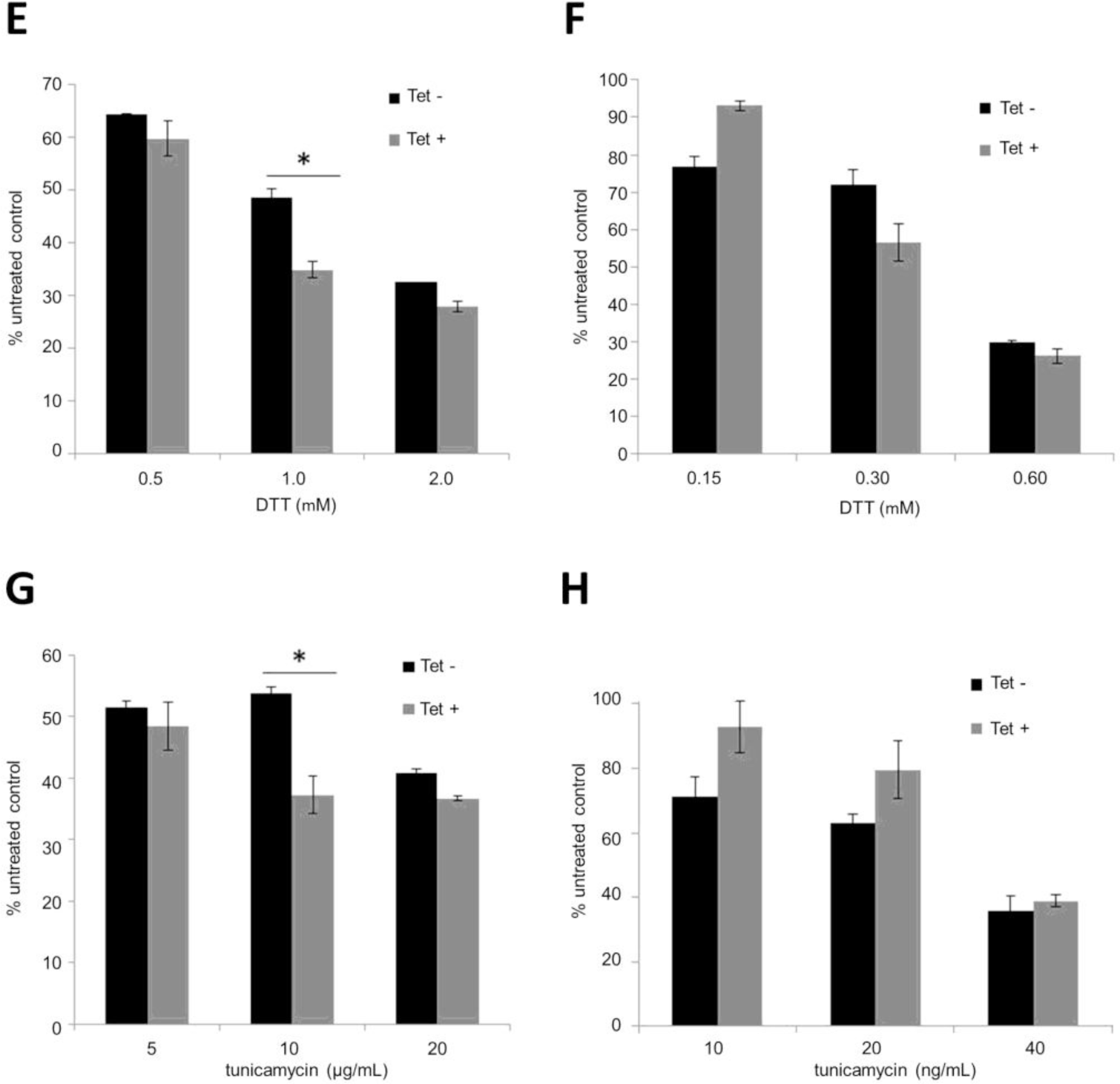
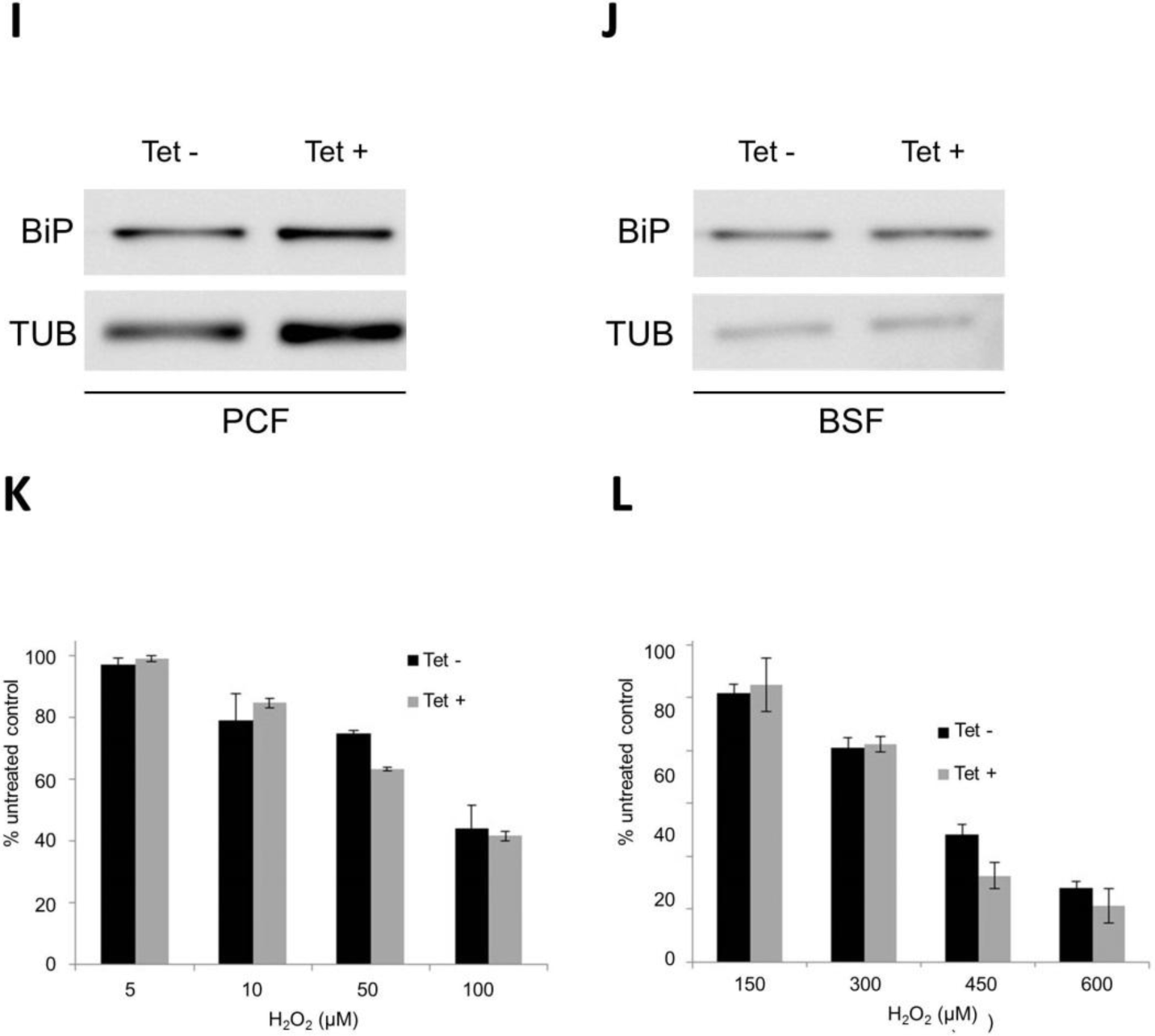
*Tb*SELENOT RNAi *T. brucei* cell lines and their sensitivity to DTT, tunicamycin and H_2_O_2_. Growth curves of *Tb*SELENOT-RNAi **A-** and **C-** PCF and **B-** and **D-** BSF *T. brucei* cell lines induced (black) and non-induced (grey) with tetracycline and real-time qPCR analysis relative to TERT as a normalization standard. Tetracycline non-induced (black) and induced (grey) *T. brucei* cells were treated with various concentrations of DTT and tunicamycin. The plots show cell concentration relative to untreated control after a 24h-incubation at 28°C and 37°C for PCF and BSF *T. brucei*, respectively. Bars represent the average of 3 independent experiments including standard deviations of experiments proceeded in **E-** and **F-** PCF *T. brucei* cells, and **G-** and **H-** BSF *T. brucei* cells. The asterisks represent significant differences between ER chemical stressors treatment (PCF *T. brucei*: 1.0 mM DTT: * P = 0.01; 10μg/mL tunicamycin: * P = 0.033; two-tailed Student’s t test). Western blot analysis of BiP in whole cell extracts of **I-** PCF and **J-** BSF *Tb*SPS2 RNAi *T. brucei* cell (12% SDS-PAGE; α-tubulin as a normalization standard). *Tb*SELENOT-RNAi **K-** PCF and **L-** BSF *T. brucei* cells were also treated with various concentrations of H_2_0_2_. Bars show the cell concentration relative to untreated controls after an 18h-incubation at 28°C and 37°C for PCF and BSF *T. brucei*, respectively. Again, the average of 3 independent experiments is shown together with the respective standard deviation.

We further evaluated the SELENOT response to ER stress upon DTT and TN treatment (Figures 6E and 6G). Interestingly, SELENOT-RNAi-induced PCF cells were more sensitive to TN than DTT (Figures 6F and 6H). On the other hand, sensitivity to different concentrations of DTT varied in SELENOT-RNAi-BSF *T. brucei,* with induced cells being more sensitive to 350-400 uM DTT. No significant effect was observed in the presence of TN (Figure 6E-H). Moreover, stable tetracycline-induced SELENOLT-RNAi lines did not show any increase in BiP expression in either PCF or BSF *T. brucei* (Figures 6I and 6J). As other selenoproteins, SELENOT contains a conserved CxxU motif [12] whose function can be related to oxidative stress defense. SEPHS2’s ability to increase trypanosome cell protection against oxidative stress caused by hydrogen peroxide [29] corroborate this statement, although it is not known whether oxidative stress protection is a consequence of selenoprotein activity or if SEPHS2 has a direct role in this process. Moreover, it is well known that ER stress increases ROS levels and vice versa in mammalian cells [47]. Therefore, we sought to test if lack of SELENOT alters the sensitivity of *T. brucei* to hydrogen peroxide. Treatment with different concentrations of hydrogen peroxide did not affect *T. brucei* growth (Figure 6J-K), ruling out a putative SELENOT role in oxidative stress protection in *T. brucei*. The contribution of other selenoproteins to oxidative stress defense awaits to be evaluated. Unfortunately, we were unable to obtain a stable SELENOK-RNAi cell line.

## Discussion

*T.brucei*, *T. cruzi* and *L. major* are known to cause severe human diseases (African trypanosomiasis, Chagas’ disease, and leishmaniasis, respectively) that mainly affect the population and economy of developing countries [1]. Besides, they are representatives of the Trypanosomatidae family that is evolutionarily distant from the most commonly studied eukaryotes (*H. sapiens*, mouse, *Drosophila melanogaster*, *Caenorhabitits elegans* and *Saccharomyces cerevisiae*) [1, 11], representing useful eukaryotic models to explore the evolution of cellular molecular processes. In fact, most of our knowledge of the eukaryotic selenocysteine pathway comes from studies of the mammalian machinery [14]. In this paper, we focused on the trypanosomatid selenophosphate synthetase, a key enzyme in the selenocysteine pathway.

The eukaryotic isoform 2 selenophosphate synthetase (SPS2, SEPHS2) is responsible for catalyzing the formation of the biological form of selenium, selenophosphate, for selenocysteine biosynthesis [14,22,38,39]. We determined the crystal structure of an N-terminally truncated selenophosphate synthetase from *L. major* (ΔN*-Lm*SEPHS2) consisting of an AIRS-like fold conserved in *E. coli* [39], *A. aeolicus* [38, 42] and *H. sapiens* [22] orthologs. This result shows that the selenophosphate synthetase fold is an important determinant for its function throughout domains of life. Although ΔN-*Lm*SEPHS2 crystallized as a monomer in the asymmetric unit, we showed that the protein is active as a dimer in solution. Our dimeric model of *Lm*SEPHS2 contains two equivalent unbound active sites in an open conformation prior to ATP and metal binding, similar to what is observed in the apo *E. coli* [39] and *A. aeolicus* [38, 42] SEPHS structures. Although partially disordered, the *Lm*SEPHS2 ATP-binding site contains conserved basic amino acid residues that lie in an extended pocket formed between the two amino acid chains of the heterodimer, explaining the need for dimerization of the full-length protein for ATPase activity. Curiously, a small amount of tetramers was also observed *in vitro* for both *L. major* and *T. brucei* SEPHS2.

A comparison between the apo selenophosphate synthetase crystal structures (*Ec*SEPHS [39] and *Aa*SEPHS [38, 42] and *Lm*SEPHS2) and the substrate-bound ones (AMPCPP-*Aa*SPS [38], ADP-*Hs*SEPHS1 [22] and AMPcP-*Hs*SEPHS1[22]) suggest that the N-terminal region of the protein, where the catalytic residues are conserved, becomes more ordered upon substrate binding. While the N-terminus of *Ec*SEPHS [39] was observed far from the open ATP-binding site, our crystal structure lacks such a flexible N-terminal region but keeps ATP-binding residues in a similar position. On the other hand, *A. aeolicus* [38] and *H. sapiens* [22] substrate-bound structures of SEPHS showed that the N-terminal region forms a long molecular tunnel suggested to preserve putative cytotoxic Se-containing intermediates from the cytoplasm.

We showed that the crystallized construct (ΔN-*Lm*SEPHS2) does not complement selenoprotein biosynthesis in SEPHS deficient *E. coli* WL400(DE3) strain, as expected due to the lack of catalytic residues. However, a residual ATPase activity was measured *in vitro* in the absence of selenide. This curious result is likely due to the preservation of the ATP-binding site being in the truncated construct. N-terminally truncated constructs of *Tb*SEPHS2 corroborate the data obtained for *Lm*SePHS2. In addition, a comparison between ΔN(25)-*Tb*SEPHS2 and ΔN(70)-*Tb*SEPHS2 functional complementation assays and ATPase activities argues that its first 25 amino acid residues are not essential for selenophosphate synthetase activity in the selenocysteine pathway, but do interfere with ATPase activity. Together, the crystallographic and functional data suggest that ATP-binding is not dependent on the N-terminal region of the protein, although ATPase activity is affected by the presence of such a region.

Interestingly, we previously reported that the disordered N-terminus of *E. coli* SEPHS is involved in the physical interaction between selenophosphate synthetase and selenocysteine synthase and is necessary for selenoprotein biosynthesis [41]. Besides, Itoh et al [38] suggested that the flexibility of the N-terminal Gly-rich loop in selenophosphate synthetase and the perselenide-carrying loop of selenocysteine lyase (SCLY) would allow the direct transfer of selenide between them. The eukaryotic selenocysteine lyase is a pyridoxal 5′-phosphate (PLP)-dependent enzyme that catalyzes the decomposition of selenocysteine into alanine and elemental selenium [19,20,48]. In fact, the homologous *E. coli* NifS-like proteins support *in vitro* selenophosphate synthesis by SEPHS in the presence of selenocysteine [49]. Moreover, *Tb*SCLY is mainly localized to the nucleus of the cell [21], while we observed that *Tb*SEPHS2 is present both in the cytoplasm and the nucleus of PCF *T. brucei* cells. However, their physical interaction had not been previously established. We demonstrate by SEC-MALS, ITC and fluorescence spectroscopy that *Tb*SEPHS2 indeed binds to *Tb*SCLY *in vitro* in the absence of tRNA^[Ser]Sec^. The SCLY active site containing PLP is obstructed in the binary complex. We further showed that both *Tb*SEPHS2 and *Tb*SCLY co-purify with each other from *T. brucei* procyclic cells and we observed that such interaction is dependent on the *Tb*SEPHS2 N-terminal region.

Furthermore, Oudouhou et al [50] also demonstrated that human SEPHS1 and SEPHS2 bind transiently to selenocysteine synthase (SEPSECS) *in vivo*. We observed that PTP-tagged selenocysteine synthase localize to the cytoplasm of PCF *T. brucei* as observed for *Tb*SEPHS2. Interestingly, *Tb*PSTK-PTP co-purified *Tb*SEPSECS and tRNA^[Ser]Sec^, demonstrating the formation of a stable ternary complex between them. However, *Tb*SEPSECS-PTP did not co-purify with any molecule, indicating that its C-terminal PTP-tag might present a steric hindrance for its interaction with *Tb*PSTK. Furthermore, neither *Tb*SEPHS2 nor *Tb*SCLY were observed in other stable complexes involved in selenocysteine biosynthesis and incorporation in selenoproteins, although a report described that selenophosphate synthetase homologs are present in higher order complexes, involved in the selenoprotein biosynthesis pathway in humans [44]. Therefore, our data do not exclude the formation of higher order transient complexes in the Sec pathway in trypanosomatids.

In addition, we showed that 80S ribosomes and polysomes involved in the synthesis of selenoproteins in PCF *T. brucei* contain *Tb*eEFSec that is dissociated in the presence of a chelating agent. A small amount of *Tb*eEFSec was also detected in the ribosome-free form. These data suggest that *Tb*eEFSec interaction with the ribosome is either dependent on selenoprotein mRNA or take place after 80S ribosome assembly or polysome association. On the other hand, *Tb*SCLY-SEPHS2 was not detected as associated to ribosomes. Similarly, *Tb*PSTK and *Tb*SEPSECS are mostly detected as ribosome-free complexes. Taken together, our biochemical data indicate that trypanosomatid selenocysteine biosynthesis occurs in a hierarchical process via coordinate action of protein complexes. We hypothesize that, after tRNA^[Ser]Sec^ aminocylation by SerRS, Ser-tRNA^[Ser]Sec^ may be either specifically transferred to a PSTK-SEPSECS binary complex or to PSTK alone, which phosphorylates Ser-tRNA^[Ser]Sec^ and subsequently associates with SEPSECS to form a stable complex. In archaea, PSTK distinguishes the characteristic D-arm of tRNA^[Ser]Sec^ over tRNA^Ser^ [51, 52] while human SEPSECS specifically recognizes its 3′-CCA end, TΨC and the variable arm [53]. Hence, Ser-tRNA^[Ser]Sec^ is discriminated from Ser-tRNA^Ser^ and mischarged tRNA^[Ser]Sec^ is avoided. Furthermore, selenophosphate is a toxic compound that is directly delivered to selenocysteine synthase by selenophosphate synthetase via a transient interaction [38, 41]. In fact, *Tb*SEPSECS was not copurified with *Tb*SEPHS2 under the conditions tested. In contrast, a stable *Tb*SEPHS2-*Tb*SCLY binary complex was obtained. A selenium delivery mechanism based on direct inter-enzyme product/substrate transfer (SCLY-SEPHS2-SEPSECS) is believed to protect the cell from selenium toxicity [38, 41]. Furthermore, neither selenophosphate synthetase nor selenocysteine synthase are specific to selenium compounds [38,54,55]. Thus, selenophosphate synthetase-selenocysteine lyase complex formation represents another level of fidelity in UGA_Sec_ codon recoding.

Curiously, selenophosphate synthetase has been described as non-essential for *T. brucei* viability [11]. SEPSECS knockout experiments further established that there is no significant contribution of selenoproteins to redox homeostasis in trypanosomatids [11,13,27,28]. Indeed, we show that *Tb*SELENOT knockdown does not significantly impact PCF and BSF *T. brucei* viability. On the other hand, our group previously demonstrated that *Tb*SEPHS2 is important for oxidative stress response in PCF *T. brucei* [29]. Accumulation of reactive oxygen species, a common characteristic of oxidative stress, can induce apoptotic cell death in *T. brucei* [56, 57]. In addition, the formation of disulfide bonds in ER proteins requires oxidizing power, which has been related to ER oxidoreductin-1 (Ero1) in mammals [58,59,60], a conserved but poorly studied protein in trypanosomatids [61]. Interestingly, protein disulfide isomerase (PDI) and thioredoxin mRNA levels increase in PCF *T. brucei* due to higher mRNA stability under DTT treatment [45]. DTT is thought to interfere with disulfide bond formation leading to accumulation of misfolded proteins in the ER [45]. Besides DTT, chemical ER stress is also commonly achieved with tunicamycin (TN), known to negatively affect N-glycosylation in the ER of *T. brucei* [62]. Here, we demonstrated that ER stress with DTT and TN upon *Tb*SEPHS2 ablation leads to growth defects in both PCF and BSF *T. brucei*, indicating a role for *Tb*SEPHS2 in the ER stress response.

Maintaining ER redox is also important for Ca^2+^-dependent cell signaling and homeostasis, which is itself key for mitochondrial homeostasis [63]. In mammals, the ER-resident selenoproteins S, N, K, M and T are believed to regulate the ER redox state, ER stress responses and Ca^2+^ signaling [64]. Interestingly, SELNOT knockout led to early rat embryonic lethality and its knockdown in corticotrope cells promoted ER stress and unfolded protein response (UPR) [65]. Among those, SELENOK and SELENOT are conserved in trypanosomatids [12, 66] and thegenome-wide tagging project in *T. brucei*, TrypTag, demonstrated that SELENOT has a reticulated cytoplasmic signal, which is compatible with endoplasmic reticulum localization [79]. Indeed, we observed that SELENOT knockdown PCF and BSF *T. brucei* cells were also sensitive to ER stress using DTT, but were not sensitive to increasing levels of hydrogen peroxide. Curiously, ER stress by TN did not lead to any negative effect to BSF *T. brucei* viability. ER function is particularly necessary for efficient production of variant surface glycoprotein (VSG) proteins that protects BSF *T. brucei* [32] surface from effectors of the host immune system [67]. Our data suggests that ER N-glycosylation of proteins is strictly regulated in trypanosomatids.

The presence of UPR is debated in *T. brucei* [32,33,35,68]. Goldshmidt et al [45] proposed that a UPR-like pathway is triggered by chemical ER stress in trypanosomatids to reduce the load of proteins to be translocated and enhance degradation of misfolded proteins. However, BiP expression was not altered upon DTT or TN treatment indicating no UPR activation as measured by an increase in BiP expression. Lack of BiP up-regulation upon chemical ER stress in *T. brucei* and *L. donovani* was previously observed by Koumandou et al [46], Izquierdo et al [69], Tiengwe et al [32] and Abhishek et al [70], arguing that a UPR-like response based on BiP is inactive in trypanosomatids. On the other hand, these parasites also conserve PKR-like endoplasmic reticulum kinase (PERK) [70], a protein that regulates protein translation by phosphorylating eIF2a, in another mechanism of UPR response in mammals [71]. It is not expected that BSF *T. brucei* could compensate for correct folding of VSGs in the absence of a UPR-like mechanism [67]. New experiments are required to more fully address the molecular response to chemical ER stressors in *T. brucei*.

The highly conserved structure of selenophosphate synthetase is essential for selenoprotein biosynthesis across domains of life. Although selenophosphate is not essential for the viability of *T. brucei* [11] and *L. donovani* [13] under laboratory controlled conditions [11], and some selenoproteins may not be essential as well, as we have shown for Selenoprotein T, our data show a role for the selenophosphate synthetase in the oxidative or ER stress protection in PCF and BSF *T. brucei*. This result is consistent with the global effect of SEPHS2 on the synthesis of selenocysteine and therefore the translation of all selenoproteins.

## Contributions

IRS and MTAS wrote this paper with help from OHT and input from all the authors. IRS and LMF prepared *Tb*/*Lm*SEPHS2 followed by native gel electrophoresis analysis with help from NKB. NKB performed HPLC-based activity assays with help from IRS and MTAS, and IRS analyzed the respective data with input from NKB and MTAS. IRS, LMF and JCB performed AUC and analyzed data. LMF solved *Tb*SEPHS2 crystal structure with help from MVBD and IRS. MLP and MTAS prepared *Tb*SCLY and performed ITC with input from IRS. MTAS performed function complementation assays in *E. coli* with help from LMF, performed RNAi and qPCR experiments with help from FCC, BM and NWA, IF with help from TCLJ and IRS, WB and polysomal profile with help from ALL, TFW, CFZ, SRV, and PTP-TAP. MTAS and IRS analyzed and interpreted the results with input from all the authors. OHT is the group leader. Funding: MTAS (FAPESP 11/24017-4 and 13/02848-7), IRS (FAPESP 10/04429-3), LMF (FAPESP 07/06591-0), FCC (FAPESP 08/58501-7), TCLJ (FAPESP 11/06087-5), OHT (FAPESP 06/55685-4, 08/57910-0). NKB, ALL and MLP are thankful for CAPES and CNPq institutional scholarships.

## Acknowledgements

We would like to acknowledge Dr. Susana Andrea Sculaccio from the São Carlos Physics Institute, University of São Paulo for technical support, Dr. Ana Carolina Migliorini Figueira from the Laboratory of Spectroscopy and Calorimetry of the National Laboratory of Biosciences (LNBIO, Brazil) for AUC, and Dr. Andressa Patricia Alves Pinto from the São Carlos Physics Institute, University of São Paulo for SEC-MALS and ITC. The *T. brucei* anti-BiP antibody was kindly provide by James D. Bangs, School of Medicine and Biomedical Sciences, State University of New York at Buffalo and trypanosome anti-EiF5a antibody by Sergio Schenkman, Departament of Microbiology, Immunology and Parasitology Federal University of São Paulo.

## Materials and Methods

### Amino-acid sequence analysis

Amino acid sequences of selenophosphate homologs were retrieved from NCBI [74]: *Aquifex aeolicus* (Aa, WP_010880640.1), *Escherichia coli* (Ec, KPO98227.1), *Pseudomonas savastanoi* (Ps, EFW86617.1), *Phytophthora infestans* (Pi, EEY58478.1), *Trypanosoma cruzi* (Tc, PBJ75389.1), *Trypanosoma brucei* (Tb, EAN78336.1), *Leishmania major* (Lm, XP_001687128.1), *Drosophila melanogaster* (Dm_1, AAB88790.1; Dm_2, NP_477478.4), *Mus musculus* (Mm_1, AAH66037.1; Mm_2, AAC53024.2), and *Homo sapiens* (Hs_1, AAH00941.1; Hs_2, AAC50958.2). Amino sequence alignment was generated using Clustal Omega [75].

### Cloning

*Tb*SEPHS2 (Tb927.10.9410), *Lm*SEPHS2 (LmjF.36.5410) and ΔN(69)*-Lm*SEPHS2 cloning was reported previously [36]. ΔN(25)-*Tb*SEPHS2, amino acid residues 26–393, and ΔN(70)-*Tb*SEPHS2, residues 71–393, were cloned into the pET20b expression vector (Novagen) using the following pairs of oligonucleotides: 5’-AGCATATGGGTCTACCGGAAGAGTTTACCTTAACTGAC-3’ and 5’-AGCTCGAGAATAATCTTATCATTTACCTTCGCTCCCACCTC-3’, and 5’-AGCATATGGATT GCAGCATTGTGAAACTGCAG-3’and 5’-AGCTCGAGAATAATCTTATCATTTACCTTCGCTCCCACCTC-3’, respectively. *Tb*SCLY (Tb927.9.12930) was cloned into the pET28b expression vector (Novagen) using the following pairs of oligonucleotides: 5’-GGATCCATGTGTAGCATTGA GGGCCCG-3’ and 5’-CTCGAGTTACTAAAACTCACCGAACTGTTGC-3’. For the PTP (Protein A-TEV site-Protein C) tagged protein constructs, the ORFs were amplified using the following primers that contained *Apa*I and *Eag*I restriction sites: *Tb*SEPHS2 5’-GGGCCCGTCTCAAATGATCCGTCCAACAG -3’ and 5’-CGGCCGAATAATCTTATCATTTACCTTC-3’, *Tb*eEFSec (Tb927.4.1820) 5’-GGGCCCCATCAC GTTTGAATGCCCTTC-3’ and 5’-CGGCCGCTGCTGAAGCTGACTGTGGAG-3’, *Tb*SEPSECS (Tb927.11.13070) 5’-GGGCC CGCCGCCATTCGACTGGGTCGTG-3’ and 5’-CGGCCGTACCCCCTCGACCGGCCAAAC-3’, *Tb*PSTK (Tb927.10.9290) 5’-CGGCCGATGACAGTTTGTCTTGTTCTAC-3’ and GGGCCCCT CGCCAAACACTTCGACTTC, *Tb*SCLY 5’-GGGCCCCTATTGATGACCTCGTGAAAC-3’ and 5’-CGGCCGAAACTCACCGAACTGTTGCAC-3’. Constructs were designed for homologous expression of C-terminally PTP-tagged protein, with exception of *Tb*PSTK, which was cloned into PN-PTP plasmid. Prior to the transfection, the constructs were linearized with the *Bsm*I, *Afl*II, *Nsi*I, *Xcm*I and *Xcm*I restriction enzymes, respectively. *Tb*SEPHS2-RNAi silencing was carried out with the construct described by Costa et al. [29] and *Tb*SELENOT-RNAi was achieved with a specific fragment PCR-amplified from PCF *T. brucei* genomic DNA using gene-specific primers 5’-CCGATTTGTTCGCATCTCATTTTC-3’ and 5’-ACCAGAGATAATTTGGCGCAG -3’ and cloned into a modified p2T7^TAblue^ with phleomycin resistance.

### Recombinant protein purification

*Tb*SEPHS2, ΔN(25)*-Tb*SEPHS2, ΔN(70)*-Tb*SEPHS2*, Lm*SEPHS2 and *ΔN-Lm*SEPHS2 were expressed in *E. coli* and purified as described previously [36]. *Tb*SCLY expression was induced by IPTG in *E. coli* BL21(DE3) for 16 hours at 18°C. Cells were harvested and sonicated in lysis buffer (50 mM de HEPES pH 7.5, 300 mM NaCl, 10 % glycerol, 10 mM imidazole, 5 mM DTT, 1X cOmplete Protease Inhibitor Cocktail (Roche)) supplemented with 10 µM PLP. The lysate was centrifuged at 40,000 X g for 20 min at 4°C and the supernatant was applied to a 5 ml Ni-NTA Superflow Cartridge (Quiagen) in ÄKTA Purifier 10 (Amersham Pharmacia Bioscience). The product was dialysed against the same buffer in the absence of imidazole, incubated with 1 mM PLP on ice and subsequently washed 5 times with the same buffer. The product was applied to a Superdex^TM^ 200 10/300 column (GE) and concentrated using an Amicon® ultracentrifugal filter.

### In vitro activity assay

Recombinant selenophosphate synthetase constructs at 30 μM final concentration were added independently to a reaction mixture (100 μl) containing 50 mM Tris-HCl pH 7.5, 50 mM KCl, 5 mM MgCl2, 5 mM DTT, 1 mM ATP and incubated at 26°C. The reaction was blocked by incubation at 75°C for 10 minutes and the solution was centrifuged at 16,000 x g for 45 minutes. The nucleotides present in the supernatant were separated by high-pressure liquid chromatography (HPLC) using a Waters Alliance 2695 HPLC with a C18 reversed-phase column (5 μm, 15 cm x 4.6 mm inside diameter, SUPELCOSIL LC-18-S; Sigma Aldrich) equipped with a guard column at a flow rate of 1 ml/min. The mobile phase consisted of buffer A (50 mM potassium phosphate buffer (KH_2_PO_4_/K_2_HPO_4_), pH 4.0) and a gradient of buffer B (20% methanol vol/vol in buffer A). The gradient conditions were: 5 min - 100% buffer A at 1.0 ml/min; 10 min - 100% buffer A at 0.1 ml/min; 1 min - 100% buffer B at 0.5 ml/min; 1 min - 100% buffer B at 1.0 ml/min; 1 min 100% buffer B at 1.0 ml/min. Nucleotide peaks were detected, and the peak area for ATP (maximum absorbance at 254 nm) was calculated relative to the respective control (enzyme absence).

### Functional complementation assay in Escherichia coli

Functional complementation experiments were conducted according to Sculaccio et al [26] for N-terminally truncated selenophosphate synthetase constructs. Briefly, *E. coli* WL400(DE3) was transformed with the full-length and truncated constructs of both *T. brucei* and *L. major* SEPHS2 and the cells were analyzed for the presence of active selenoprotein formate dehydrogenase H (FDH H) using the benzyl viologen assay under anaerobic conditions at 30°C for 48 hours.

### Native gel electrophoresis

Recombinant *Tb*SEPHS2, ΔN(25)*-Tb*SEPHS2, ΔN(70)*-Tb*SEPHS2*, Lm*SEPHS2 and *ΔN-Lm*SEPHS2 were applied to a PhastGel gradient 8-25% (GE) at 1 mg/ml at room temperature.

### Analytical ultracentrifugation (AUC)

*Tb*SEPHS2 and *Lm*SEPSH2 at 0.15, 0.30, 0.45, 0.60 and 0.80 mg/ml in 25 mM Tris pH 8.0, 50 mM NaCl, 1mM β-mercaptoethanol were subjected to velocity sedimentation at 30,000 rpm at 20°C in an An60Ti rotor using a OptimaTM XL-A analytical ultracentrifuge (Beckmann). The data were analyzed with SEDFIT using a c(s) distribution model. The partial-specific volumes (v-bar), solvent density and viscosity were calculated using SEDNTERP (Dr Thomas Laue, University of New Hampshire). To determine the tetramer-dimer equilibrium dissociation, 110 µL of protein at a concentrations of 0.5, 0.75 and 1.0 mg/ml were loaded in 12 mm 6-sector cells and centrifuged at 8,000, 10,000 and 12,000 rpm at 4°C until equilibrium had been reached. Data were processed and analyzed using SEDPHAT.

### Size exclusion chromatography with multi angle light scattering (SEC-MALS)

The molecular mass distribution of *Tb*SCLY (40 μM), *Tb* SEPHS2 (40 μM) and *Tb-SCLY-SEPHS2* (40 μM:40 μM) in solution was determined using SEC-MALS. A 1:1 *Tb-SCLY-* SEPHS2 mixture was incubated in 50 mM de HEPES pH 7.5, 300 mM NaCl, 10 % glycerol, 10 mM imidazole, 5 mM DTT at 25°C for 30 minutes and 60 minutes on ice previous to the SEC-MALS analysis. 40 μl of each sample was loaded onto a WTC-030N5 (Wyatt) column running at 0.3 ml/min coupled to a mini-DAWN® TREOS® equipped with an Optilab® rEX detector (Wyatt). Data was analyzed using Astra 7.0.1.24 (Wyatt). Experiments were performed at room temperature.

### Isothermal titration calorimetry (ITC)

ITC measurements were performed at 25°C in a VP-ITC calorimeter (Microcal) using 50 mM Tris-Cl, pH 7.4, 100 mM NaCl, 1 mM DTT buffer. Samples of *Tb*SCLY at 10 nM in the calorimeter cell were titrated with *Tb*SEPHS2 at 200 μM from the syringe using 29 injections of 2 μL preceded by a 0.5 μL injection. The resulting excess heats associated with the injections were integrated and normalized using the measured concentrations of protein from UV absorbance, corrected for the background heat of dilution of *Tb*SEPHS2 and the binding data were fitted to a two-species hetero-association model using Microcal ORIGIN software (Microcal). We were unable to perform the potentially informative reverse titration, with *Tb*SCLY in the syringe, because of its limited stability at high concentration at 25°C.

### Fluorescence spectroscopy

Fluorescence spectroscopy of the pyridoxal phosphate (PLP) group bound to *TbSCLY* excited at 450 nm was performed in an ISS-PC spectrofluorometer (ISS). Varying concentrations of *Tb*SEPHS2 (1:0.5, 1:0.75, 1:1, 1:1.25, 1:1.5) were incubated in 50 mM de HEPES pH 7.5, 300 mM NaCl, 10 % glycerol, 10 mM imidazole, 5 mM DTT with 20 μM *Tb*SCLY at 25°C for 30 minutes and 60 minutes on ice. Fluorescence spectroscopy was then performed at room temperature.

### Circular Dichroism (CD) Spectroscopy

Purified *Tb*SEPHS2, *Lm*SEPHS2 and their truncated constructs were dialyzed against 25 mM Tris pH 8.0, 50 mM NaCl, 1mM de β-mercaptoethanol at 4°C overnight and adjusted to a concentration of 0.2 mg/ml. Purified *Tb*SCLY at 0.2 mg/ml was kept in the same buffer. Far-UV CD spectra at 5°C were measured using a Jasco J-815 spectropolarimeter (JASCO) (0,1 nm resolution, 100 nm/min, quartz 0.5 cm cuvette).

### X-ray crystallography

X-ray diffraction data collection and analysis were previously reported for both *Tb*SEPHS2 and ΔN-*Lm*SEPHS2 crystals. Although the data was not sufficient for *Tb*SEPHS2 structure determination, ΔN-*Lm*SEPHS2 final model was completed by molecular replacement with PHASER [76] using *H. sapiens* SEPHS1 [22] structure coordinates as search model (PDB code 3FD5). Model building and refinement were performed using PHENIX [77] and COOT [78]. Structure visualization was performed in PyMOL. ConSurf [72] was used for conservation analysis.

### RNAi experiments

*Trypanosoma brucei* procyclic RNAi host strain 29-13 was grown in SM-9 [80] supplemented with 10% FBS tetracycline-tested (Atlanta Biologicals) in the presence of G418 (15 μg**/**ml) and hygromycin (50 μg**/**ml) to maintain the integrated genes for T7 RNA polymerase and tetracycline repressor, respectively. *T. brucei* bloodstream form, strain 221, was cultured in HMI-9 media supplemented with 10% FBS tetracycline-tested, supplemented with G418 (15 μg**/**ml).

The RNAi constructs were obtained according to Costa et al. 2011. Transfected cells were cloned by limiting dilutions and selected with 2.5 μg/mL of phleomycin (Sigma).To monitor the growth of RNAi cells, dsRNA synthesis was induced with 1 μg/mL of tetracycline and the cells were counted daily and diluted to a concentration of 2 x10^5^ cells/mL.

For bloodstream transfection it was used the protocol described by Burkard and coworkers [81]. In sum, 10 μg of *Not*I-linearized DNA were used per 6×10^7^ cells in 100μl homemade Tb-BSF buffer (90mM sodium phosphate, 5mM potassium chloride, 0.15mM calcium chloride, 50mM HEPES, pH 7.3). Electroporation was performed using 2mm gap cuvettes (BTX, Harvard apparatus) with program X-001 of the Amaxa Nucleofector (Lonza). Following each transfection, stable transformants were selected for 6 days with 2.5 μg/ml pheomycin as a pool culture in 125 ml HMI-9 medium containing 10% fetal bovine serum.

Mid-log phase *T. brucei* procyclic form (5 x 10^5^ cells/mL) and *T. brucei* bloodstream form (1 x 10^5^ cells/mL) were treated with tetracycline for 24 hours aimed to induced the RNAi. These cells were incubated with various DTT and tunicamycin concentrations for 4 hours or H_2_O_2_ for 8 hours and cell viability was confirmed by staining with fluorescein diacetate (FDA, Sacks and Melby, 1998).

### Real Time RT-PCR (qPCR)

RNA from 1 to 2×10^7^ trypanosomes bloodstream and procyclic forms, respectively, were isolated with NucleoSpin RNA II Kit, (Macherey-Nagel), and 1.2 μg of RNA were reverse transcribed by SuperScript III Reverse Transcriptase (Life Technologies). To quantify levels of specific mRNA transcripts in individual samples, 1 μl of each cDNA sample was amplified with gene-specific primers in iQ SYBR Green Supermix (Bio-Rad) according to the manufacturer’s protocol, using a C1000 thermocycler fitted with a CFX96 real-time system (Bio-Rad Laboratories). Three technical and three biological replicates of each reaction were performed. Every test gene was normalized to the mRNAs encoding TERT, which is known to be expressed constitutively (Brenndörfer and Boshart, 2011). The temperature profile was: 95°C, 3min [(95°C, 15 s; 55.5°C, 1 min; data collection) 40°C].

### Polysomal profile analysis

For polysome profile analysis, 500 mL culture of each *T. brucei* PTP lineage were grown overnight at 28°C to mid-log phase and cells were treated with cyclohexamide 100 μg/mL for 5 minutes. The cultures were immediately chilled on ice and collected by centrifugation at 3,000 xg at 4°C for 7 minutes. Cells were washed twice with ice-cold Salts buffer (Tris-HCl 10 mM pH 7,5; KCl 30 mM; MgCl_2_; 10 mM; DTT 1 mM; cyclohexamide 100 μg/mL), pellet volume was estimated and suspended with the same volume of Lysis buffer (Tris-HCl 10 mM pH 7,5; KCl 30 mM; MgCl_2_; 10 mM; DTT 1 mM; cyclohexamide 100 μg/mL; 1,2% Triton), supplemented with 1X Protease cOmplete inhibitor cocktail (Roche). Lysates were clarified by centrifugation at 17,000 xg for 15 minutes. Eight hundred micrograms equivalent of OD_260_ units was loaded on a 7–47% sucrose gradient prepared in Salts buffer and centrifuged at 39,000 RPM for 2 hours at 4°C in a Beckman SW41Ti rotor. The gradients were fractionated by upward displacement with 60% (w/v) sucrose using a gradient fractionator ISCO UA-6 UV Vis with Type 11 optical unit at 254 nm and fractions were collected manually for subsequent western blotting analysis. Subunit profile analysis was performed as described before with modifications; lysis buffer was supplemented with 50 mM of EDTA and lysed was centrifuged on a 5–25% sucrose gradient for 4 hours.

### PTP-tag TAP and Mass Spectrometry

PTP-tagged protein expression was demonstrated by immunoblotting with a rabbit polyclonal anti-protein A antibody (SIGMA) as described previously [13]. Tandem affinity purification (TAP) was performed according to a standard PTP purification protocol [42].

For protein identification, bands were excised from the gel, de-stained with 50% methanol and 5% acetic acid for 5 minutes, dehydrated with 100% acetonitrile for another 5 minutes and reduced with 5 mM DTT for 30 min at room temperature, following alkylation with 14 mM of iodoacetamide in the dark for 30 min. Proteins digest were carried out with 0.75 µg of trypsin (SIGMA) for 16 h at 4°C, under agitation (900 rpm) and reactions were stopped with 5% formic acid and bands dried in vacuum. Digestion products were desalted using ZipTipC18 (Merck) according the manufacturer’s instructions. Peptides were suspended in 0.1% formic acid and injected in an in-house made 5 cm reversed phase pre-column (inner diameter 100 µm, filled with a 10 µm C18 Jupiter resins-Phenomenex) coupled to a nano-HPLC (NanoLC-1DPlus, Proxeon) online to an LTQ-Orbitrap Velos (ThermoFisher Scientific). The peptide fractionation was carried out on an in-house 10 cm reversed phase capillary emitter column (inner diameter 75 µm, filled with 5 µm C18 Aqua resins-Phenomenex) with a gradient of 2-35 % of acetonitrile in 0.1% formic acid for 52 min followed by a gradient of 35-95% for 5 min at a flow rate of 300 ml/min. The mass spectrometry was operated in a data-dependent acquisition mode at 1.9 kV and 200°C. MS/MS spectra were acquired at normalized collision energy of 35% with FT scans from m/z 200 to 2000 and mass resolution of 3 kDa. Raw data were processed in Proteome Discovery 1.3 using MASCOT as a search engine and the complete database of *T. brucei* obtained from TriTrypDB [32].

### RNA analysis

Reverse transcription (RT)-PCR experiments were used to monitor the presence of tRNA^[Ser]Sec^ in selenocysteine protein complexes. Protein pull-down assays, 100 μl cell extracts were incubated with 30 μl of IgG sepharose 6 fast flow (GE) beads, equilibrated with PA-150 buffer. After five washes, total RNA was extracted by TRIzol reagent (GE) and the first strand synthesized by SuperScript III reverse transcriptase (Invitrogen) with random hexamers primers (ThermoFisher scientific). PCR was performed with tRNA^[Ser]Sec^ sense (5’ GCGCCACGATGAGCTCAGCTG 3’) and tRNA^[Ser]Sec^ antisense (5’ CACCACAAAGGCCGAATCGAAC 3’) oligonucleotides.

### Fluorescence microscopy

*T. brucei* cell lines expressing PTP-tagged proteins were used for immunolocalization assays. Briefly, 5 × 10^6^ cells were washed with PBS buffer pH 7.4 (SIGMA), fixed with 2% v/v paraformaldehyde for 20 min at 4°C, permeabilized with 0.3% v/v Triton X-100 for 3 min at 4°C. Cells were blocked with 3% w/v BSA and incubated with 1:16,000 v/v rabbit anti-protein A antibody (SIGMA) for 1 h at room temperature. After washes, cells were incubated with 1:400 (v/v) Alexa Fluor® 594-conjugated (Invitrogen) and 10 μg/ml DAPI (SIGMA) at room temperature. The coverslips were mounted on glass slides with Vectashield mounting medium (Vector Laboratories) and images obtained with Olympus IX-71 (Olympus) inverted microscope coupled Photometrix CoolSnapHQ CCD camera were further deconvoluted using DeltaVision (Applied Precision) software.

## Supplementary Information

**Table S1:**
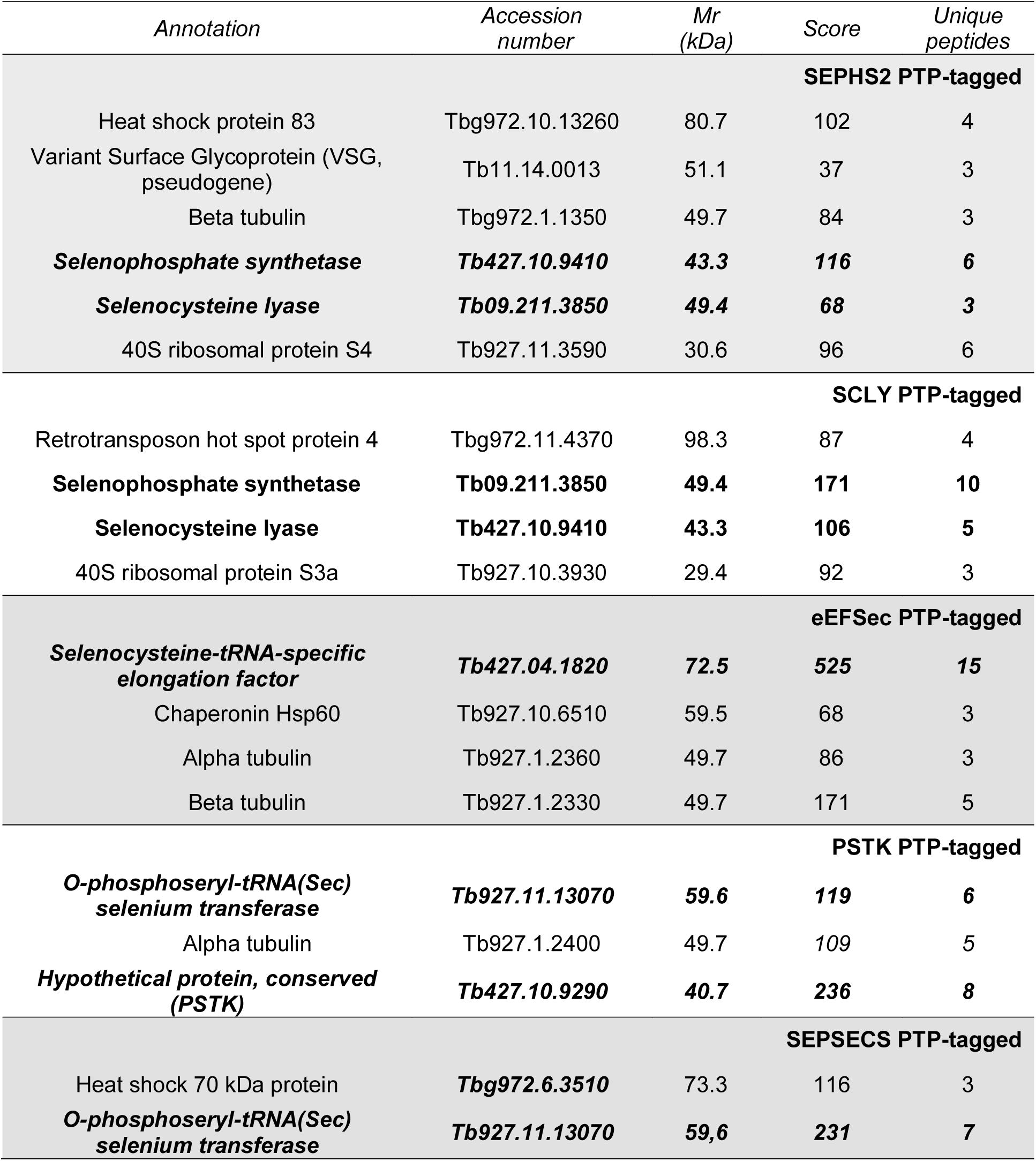
Mass spectrometry identification of co-eluted proteins with selenocysteine machinery components.

**Figure S1:**
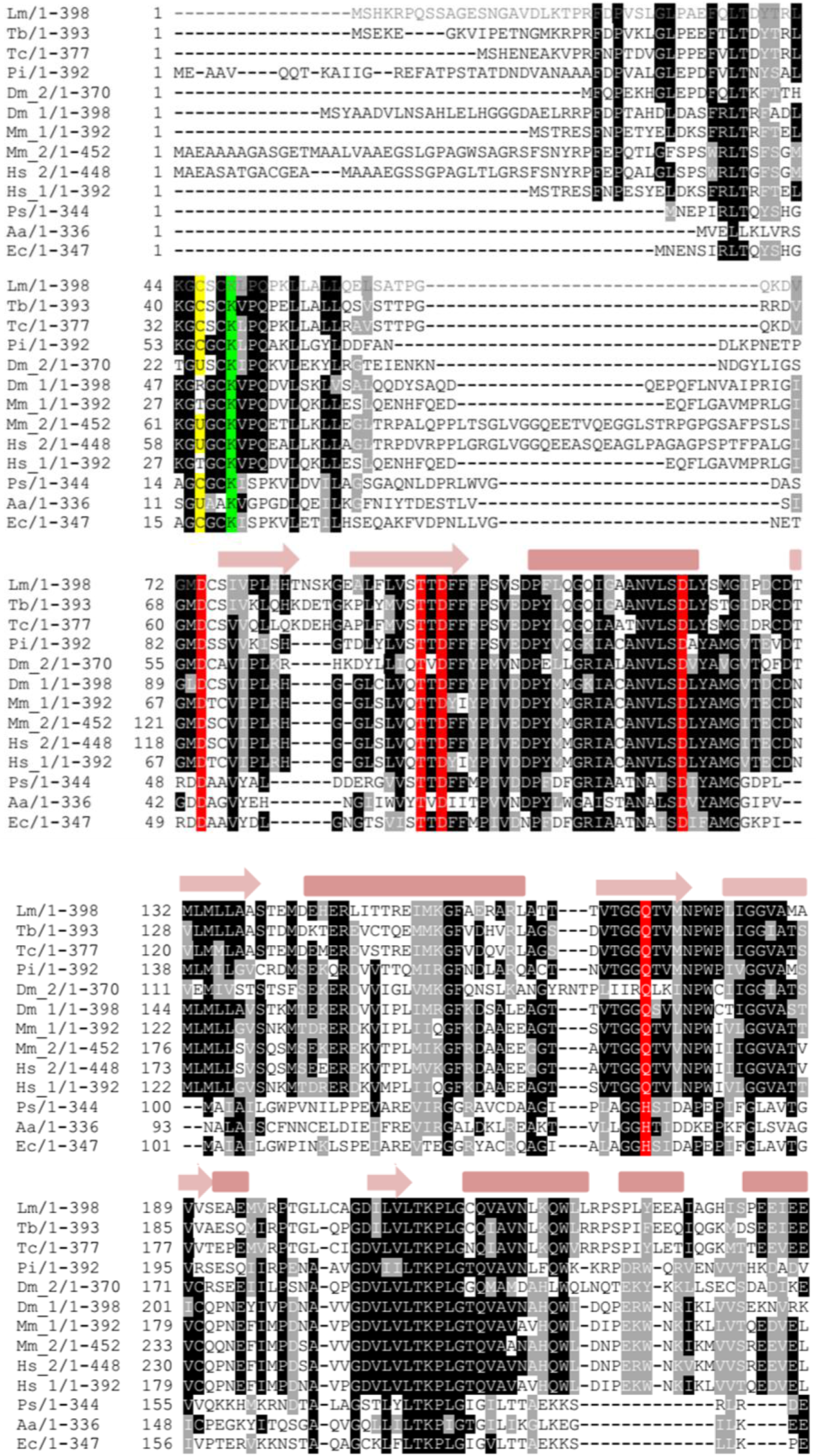
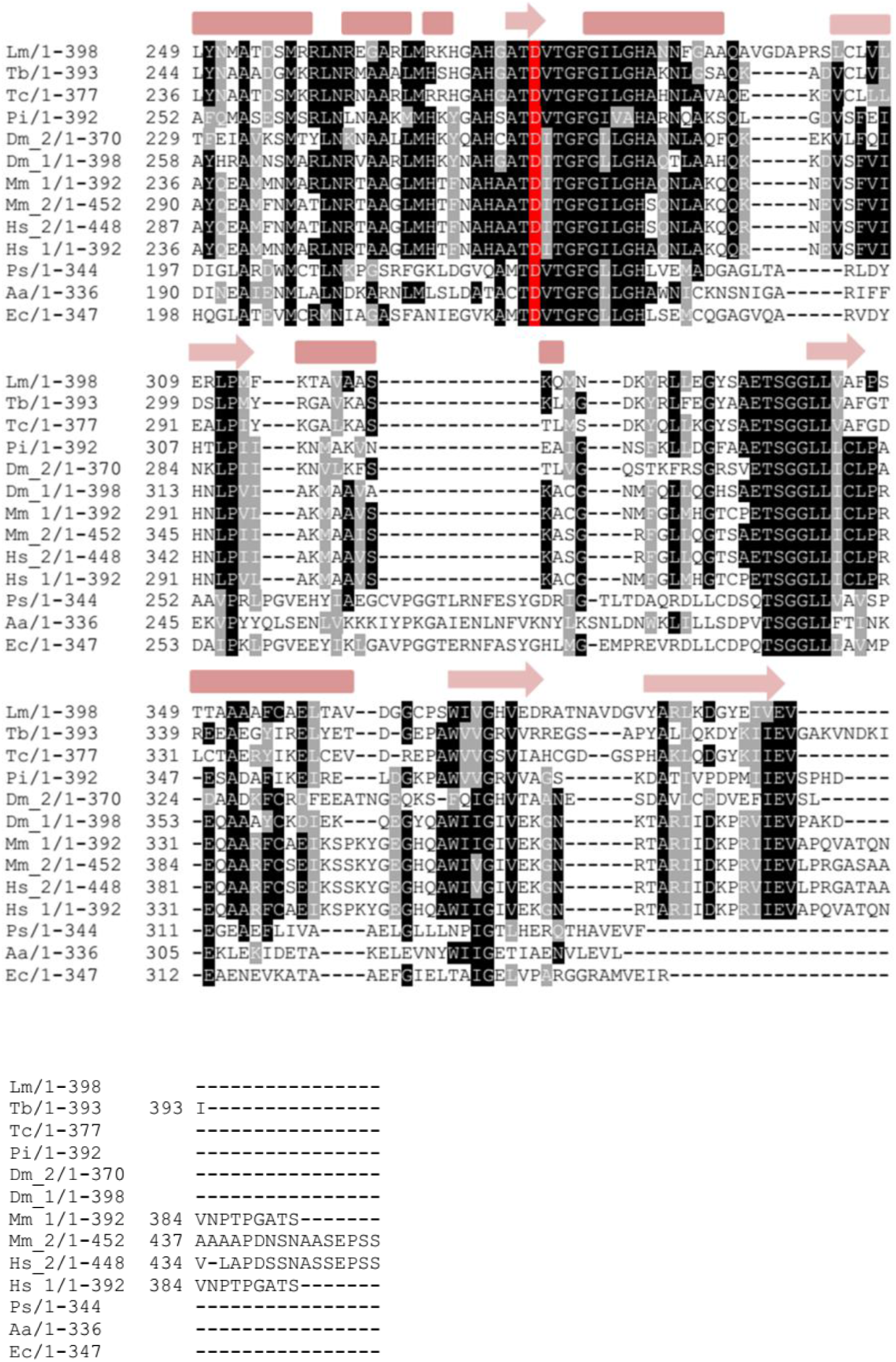
Multiple amino acid sequence alignment. Conserved residues are highlighted (catalytic Cys/Sec in yellow and Lys in green, and ATP binding amino acids in red). Amino acid sequences: Lm – *L. major* (XP_001687128.1), Tb – *T. brucei* (EAN78336.1), Tc – *T. cruzi* (PBJ75389.1), Pi – *Phytophthora infestans* (EEY58478.1), Dm - *Drosophila melanogaster* (Dm_1: AAB88790.1, Dm_2: NP_477478.4), Mm - *Mus musculus* (Mm_1: AAH66037.1, Mm_2: AAC53024.2), Hs – *Homo sapiens* (Hs_1: AAH00941.1, Hs_2: AAC50958.2), Ps - *Pseudomonas savastanoi* (EFW86617.1), Aa – *Aquifex aeolicus* (WP_010880640.1), Ec - *Escherichia coli* (KPO98227.1).

**Figure S2:**
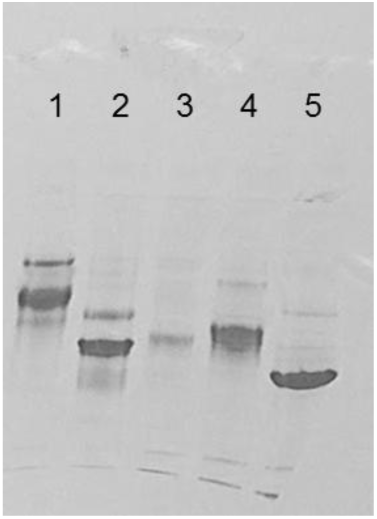
Native gel electrophoresis of selenophosphate synthetase constructs. 1- *Tb*SEPHS2, 2- ΔN(70)-*Tb*SEPHS2, 3- ΔN(25)-*Tb*SEPHS2, 4- *Lm*SEPHS2, and 5- ΔN(69)-*Lm*SEPHS2. Major bands correspond to dimers. The second most abundant species in each lane corresponds to the respective tetramer.

**Figure S3:**
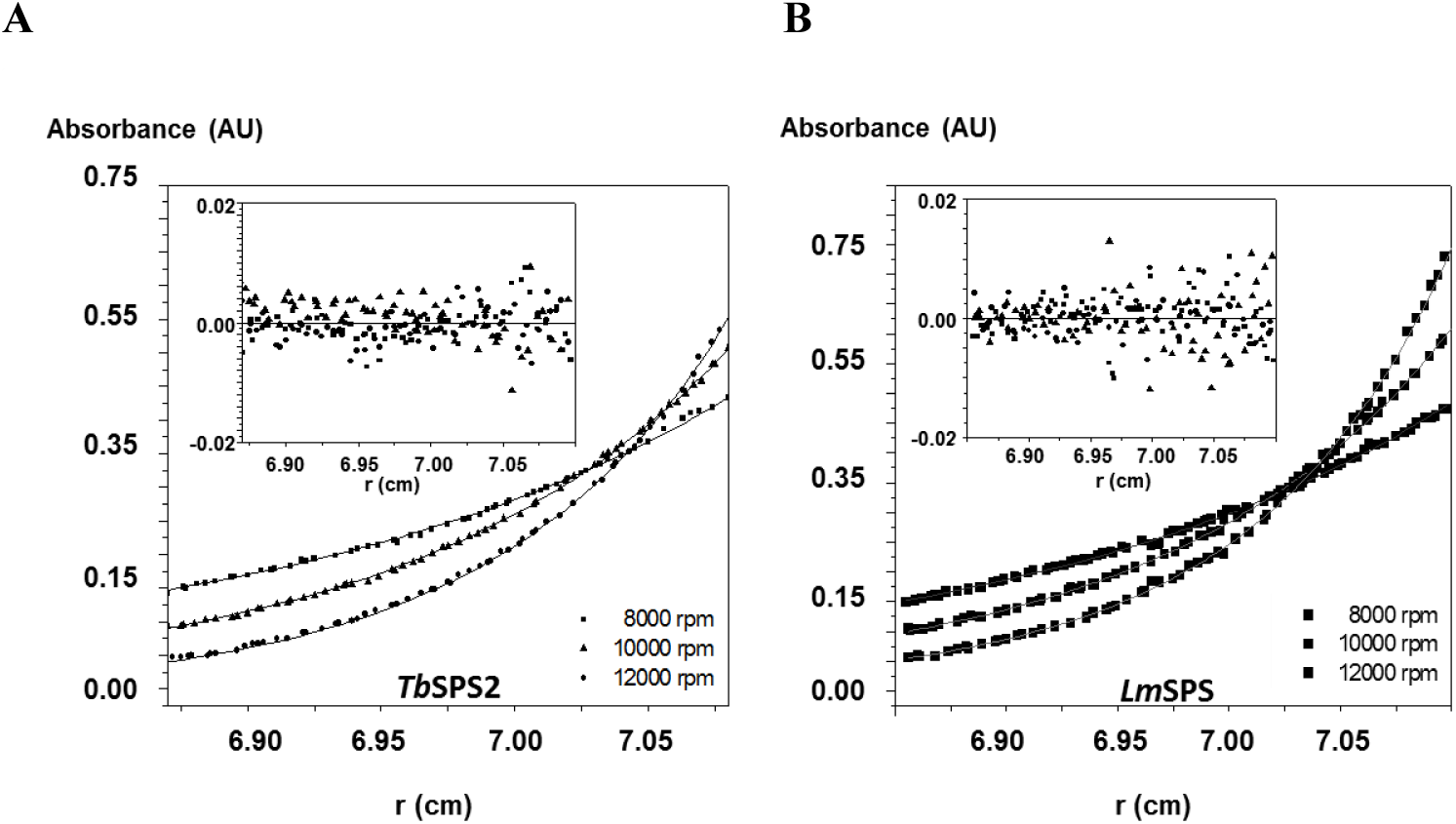
Sedimentation equilibrium analytical ultracentrifugation (SE-AUC). Multi-speed and multi-concentration global fitting (SEDPHAT [79]) for a dimer-tetramer self-association system of A- *Tb*SEPHS2 (K_d_ = 161 ± 10 μM) and B- *Lm*SEPHS2 (K_d_ = 178 ± 10 μM). Fitting residuals are shown as inserts.

**Figure S4:**
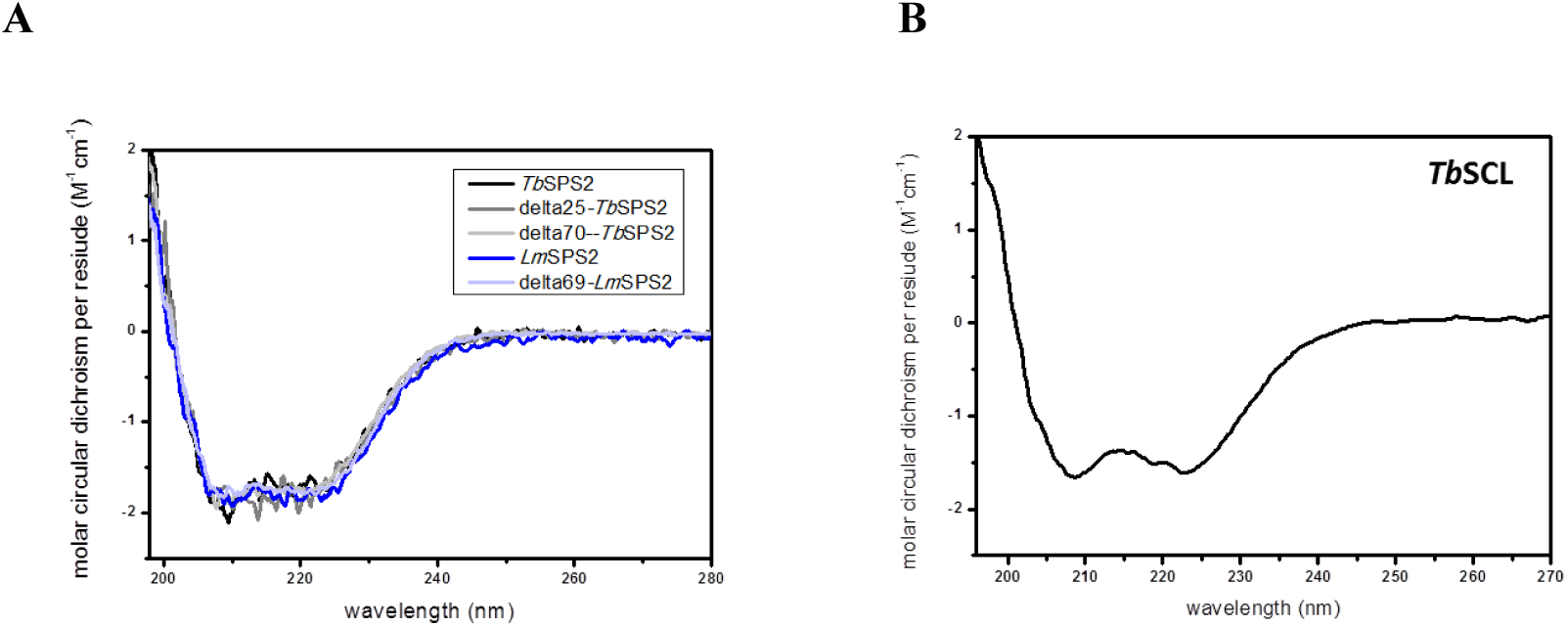
Circular dichroism (CD) spectroscopy. CD spectra for A- selenophosphate synthetase constructs (*Tb*SEPHS2, ΔN(25)-*Tb*SEPHS2, ΔN(70)-*Tb*SEPHS2, *Lm*SEPHS2 and ΔN-*Lm*SEPHS2, and B- *T. brucei* selenocysteine lyase (*Tb*SCLY).

**Figure S5:**
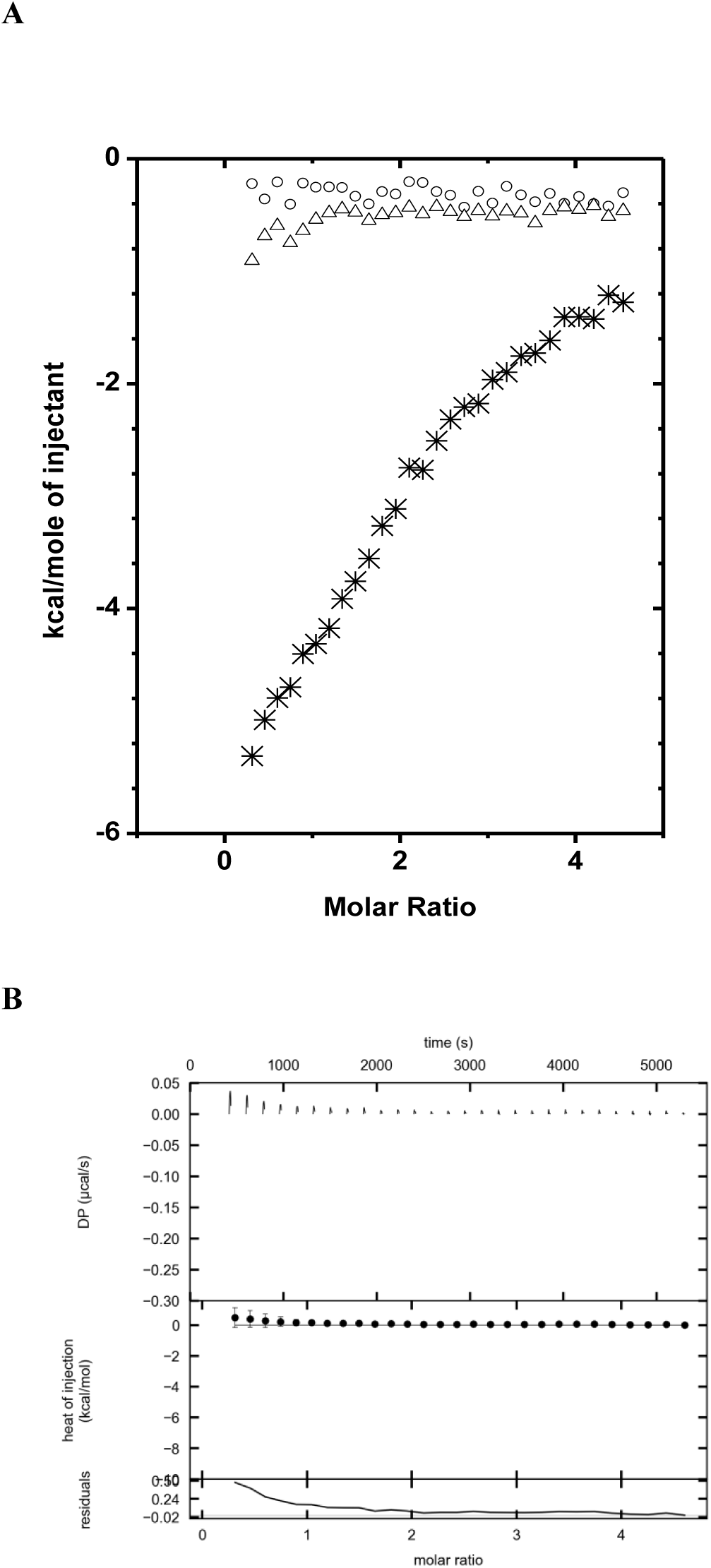
ITC. A- ITC data for SCLY-buffer (circle), buffer-SEPHS2 (triangle) and SEPHS2-SCLY (star), and B-SCLY-ΔN(70)-SEPHS2 titration experiments using VP-ITC calorimeter and analyzed in NITPIC.

**Figure S6:**
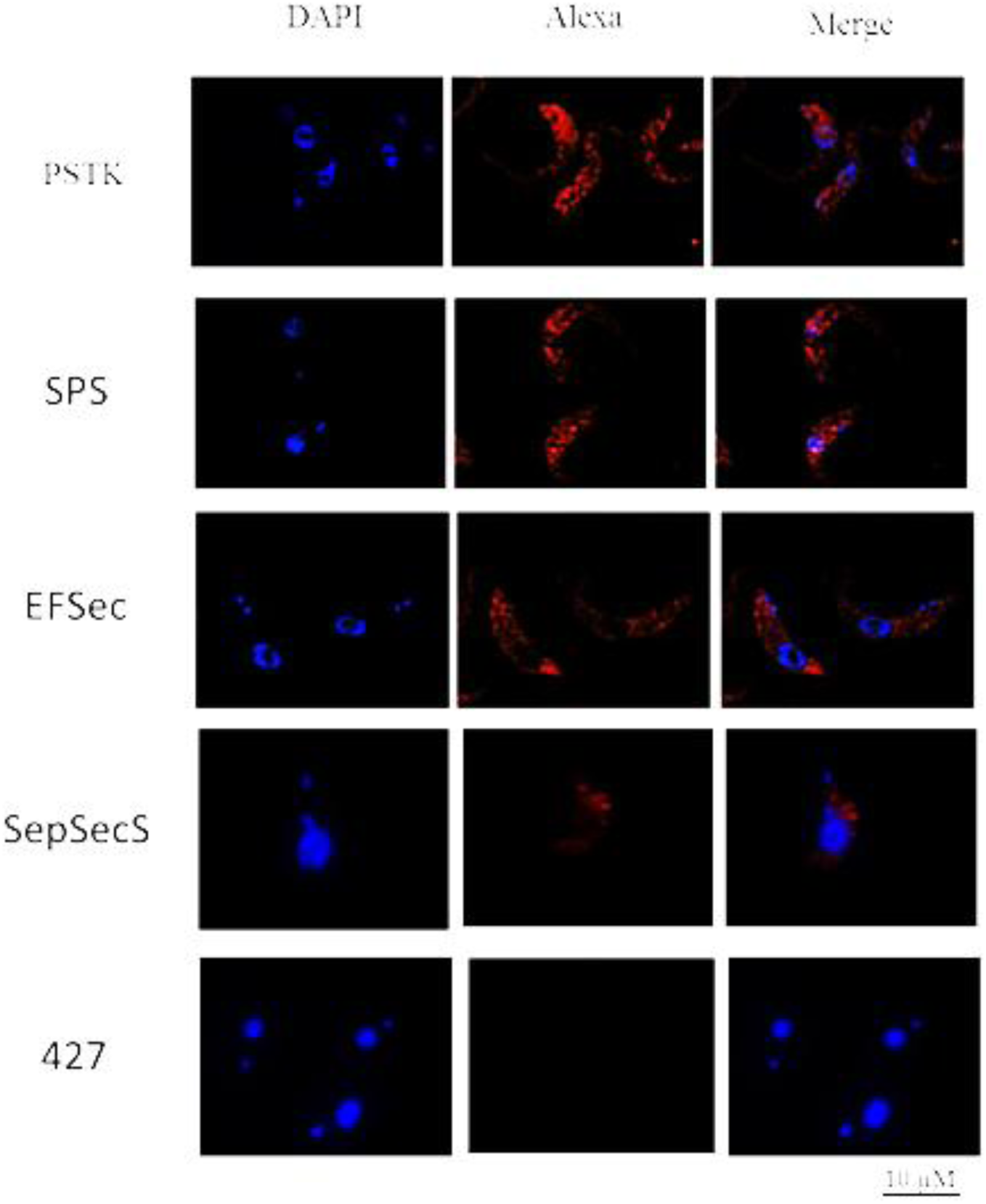
Selenocysteine pathway machinery localization in procyclic *T. brucei* cells. PTP-tagged proteins immunolocalized using anti-protein A antibody (red). DAPI (blue) is used as a nuclear/kinetoplast marker. Untransfected *procyclic T. brucei* 427 cells were used as negative controls.

## References

1. Kaufer A, Ellis J, Stark D, Barratt J. The evolution of trypanosomatid taxonomy. Parasites & Vectors. 2017;10. doi: 10.1186/s13071-017-2204-7. PubMed PMID: WOS:000402847200001.

2. Franco JR, Simarro PP, Diarra A, Jannin JG. Epidemiology of human African trypanosomiasis. Clinical epidemiology. 2014;6:257–75. doi: 10.2147/clep.s39728. PubMed PMID: MEDLINE:25125985.

3. Messenger LA, Miles MA, Bern C. Between a bug and a hard place: Trypanosoma cruzi genetic diversity and the clinical outcomes of Chagas disease. Expert review of anti-infective therapy. 2015;13(8):995–1029. doi: 10.1586/14787210.2015.1056158. PubMed PMID: MEDLINE:26162928.

4. Kevric I, Cappel MA, Keeling JH. New World and Old World Leishmania Infections: A Practical Review. Dermatologic clinics. 2015;33(3):579–93. doi: 10.1016/j.det.2015.03.018. PubMed PMID: MEDLINE:26143433.

5. Daniels J-P, Gull K, Wickstead B. Cell Biology of the Trypanosome Genome. Microbiology and Molecular Biology Reviews. 2010;74(4):552–69. doi: 10.1128/mmbr.00024-10. PubMed PMID: WOS:000284758500004.

6. Guenzl A. The Pre-mRNA Splicing Machinery of Trypanosomes: Complex or Simplified? Eukaryotic Cell. 2010;9(8). doi: 10.1128/ec.00113-10. PubMed PMID: WOS:000280577300003.

7. Fairlamb AH, Cerami A. METABOLISM AND FUNCTIONS OF TRYPANOTHIONE IN THE KINETOPLASTIDA. Annual Review of Microbiology. 1992;46:695–729. doi: 10.1146/annurev.micro.46.1.695. PubMed PMID: WOS:A1992JQ92900024.

8. Krauth-Siegel RL, Comini MA. Redox control in trypanosomatids, parasitic protozoa with trypanothione-based thiol metabolism. Biochimica Et Biophysica Acta-General Subjects. 2008;1780(11):1236–48. doi: 10.1016/j.bbagen.2008.03.006. PubMed PMID: WOS:000259359600005.

9. Manta B, Comini M, Medeiros A, Hugo M, Trujillo M, Radi R. Trypanothione: A unique bis-glutathionyl derivative in trypanosomatids. Biochimica Et Biophysica Acta-General Subjects. 2013;1830(5):3199–216. doi: 10.1016/j.bbagen.2013.01.013. PubMed PMID: WOS:000317797600008.

10. Steinbrenner H, Speckmann B, Klotz LO. Selenoproteins: Antioxidant selenoenzymes and beyond. Archives of Biochemistry and Biophysics. 2016;595:113–9. doi: 10.1016/j.abb.2015.06.024. PubMed PMID: WOS:000374705000023.

11. Aeby E, Palioura S, Pusnik M, Marazzi J, Lieberman A, Ullu E, et al. The canonical pathway for selenocysteine insertion is dispensable in Trypanosomes. Proceedings of the National Academy of Sciences of the United States of America. 2009;106(13):5088–92. doi: 10.1073/pnas.0901575106. PubMed PMID: WOS:000264790600026.

12. Lobanov AV, Gromer S, Salinas G, Gladyshev VN. Selenium metabolism in Trypanosoma: characterization of selenoproteomes and identification of a Kinetoplastida-specific selenoprotein. Nucleic Acids Research. 2006;34(14):4012–24. doi: 10.1093/nar/gkl541. PubMed PMID: WOS:000240583800022.

13. Manhas R, Gowri VS, Madhubala R. Leishmania donovani Encodes a Functional Selenocysteinyl-tRNA Synthase. Journal of Biological Chemistry. 2016;291(3):1203–20. doi: 10.1074/jbc.M115.695007. PubMed PMID: WOS:000368068300017.

14. Serrao VHB, Silva IR, da Silva MTA, Scortecci JF, Fernandes AD, Thiemann OH. The unique tRNA(Sec) and its role in selenocysteine biosynthesis. Amino Acids. 2018;50(9):1145–67. doi: 10.1007/s00726-018-2595-6. PubMed PMID: WOS:000442584500001.

15. Xu X-M, Carlson BA, Mix H, Zhang Y, Saira K, Glass RS, et al. Biosynthesis of selenocysteine on its tRNA in eukaryotes. Plos Biology. 2007;5(1):96–105. doi: 10.1371/journal.pbio.005004. PubMed PMID: WOS:000245243100010.

16. Bulteau AL, Chavatte L. Update on Selenoprotein Biosynthesis. Antioxidants & Redox Signaling. 2015;23(10):775–94. doi: 10.1089/ars.2015.6391. PubMed PMID: WOS:000361750400003.

17. da Silva MTA, Silva-Jardim I, Thiemann OH. Biological Implications of Selenium and its Role in Trypanosomiasis Treatment. Current Medicinal Chemistry. 2014;21(15):1772–80. PubMed PMID: WOS:000334356600008.

18. Allmang C, Wurth L, Krol A. The selenium to selenoprotein pathway in eukaryotes: More molecular partners than anticipated. Biochimica Et Biophysica Acta-General Subjects. 2009;1790(11):1415–23. doi: 10.1016/j.bbagen.2009.03.003. PubMed PMID: WOS:000271454600004.

19. Collins R, Johansson A-L, Karlberg T, Markova N, van den Berg S, Olesen K, et al. Biochemical Discrimination between Selenium and Sulfur 1: A Single Residue Provides Selenium Specificity to Human Selenocysteine Lyase. Plos One. 2012;7(1). doi: 10.1371/journal.pone.0030581. PubMed PMID: WOS:000301640600038.

20. Tobe R, Mihara H. Delivery of selenium to selenophosphate synthetase for selenoprotein biosynthesis. Biochimica Et Biophysica Acta-General Subjects. 2018;1862(11):2433–40. doi: 10.1016/j.bbagen.2018.05.023. PubMed PMID: WOS:000444931900012.

21. Poliak P, Van Hoewyk D, Obornik M, Zikova A, Stuart KD, Tachezy J, et al. Functions and cellular localization of cysteine desulfurase and selenocysteine lyase in Trypanosoma brucei. Febs Journal. 2010;277(2):383–93. doi: 10.1111/j.1742-4658.2009.07489.x. PubMed PMID: WOS:000273396000013.

22. Wang KT, Wang J, Li LF, Su XD. Crystal Structures of Catalytic Intermediates of Human Selenophosphate Synthetase 1. Journal of Molecular Biology. 2009;390(4):747–59. doi: 10.1016/j.jmb.2009.05.032. PubMed PMID: WOS:000268519200014.

23. Lobanov AV, Hatfield DL, Gladyshev VN. Selenoproteinless animals: Selenophosphate synthetase SPS1 functions in a pathway unrelated to selenocysteine biosynthesis. Protein Science. 2008;17(1):176–82. doi: 10.1110/ps.073261508. PubMed PMID: WOS:000251834500020.

24. Xu XM, Carlson BA, Irons R, Mix H, Zhong NX, Gladyshev VN, et al. Selenophosphate synthetase 2 is essential for selenoprotein biosynthesis. Biochemical Journal. 2007;404:115–20. doi: 10.1042/bj20070165. PubMed PMID: WOS:000246547300013.

25. Na J, Jung J, Bang J, Lu Q, Carlson BA, Guo X, et al. Selenophosphate synthetase 1 and its role in redox homeostasis, defense and proliferation. Free Radical Biology and Medicine. 2018;127:190–7. doi: 10.1016/j.freeradbiomed.2018.04.577. PubMed PMID: WOS:000445890700021.

26. Sculaccio SA, Rodrigues EM, Cordeiro AT, Magalhaes A, Braga AL, Alberto EE, et al. Selenocysteine incorporation in Kinetoplastid: Selenophosphate synthetase (SELD) from Leishmania major and Trypanosoma brucei. Molecular and Biochemical Parasitology. 2008;162(2):165–71. doi: 10.1016/j.molbiopara.2008.08.009. PubMed PMID: WOS:000261164700008.

27. Aeby E, Seidel V, Schneider A. The selenoproteome is dispensable in bloodstream forms of Trypanosoma brucei. Molecular and Biochemical Parasitology. 2009;168(2):191–3. doi: 10.1016/j.molbiopara.2009.08.007. PubMed PMID: WOS:000271126900010.

28. Bonilla M, Krull E, Irigoin F, Salinas G, Comini MA. Selenoproteins of African trypanosomes are dispensable for parasite survival in a mammalian host. Molecular and Biochemical Parasitology. 2016;206(1-2):13–9. doi: 10.1016/j.molbiopara.2016.03.002. PubMed PMID: WOS:000379373600003.

29. Costa FC, Oliva MAV, de Jesus TCL, Schenkman S, Thiemann OH. Oxidative stress protection of Trypanosomes requires selenophosphate synthase. Molecular and Biochemical Parasitology. 2011;180(1):47–50. doi: 10.1016/j.molbiopara.2011.04.007. PubMed PMID: WOS:000295545000008.

30. Pitts MW, Hoffmann PR. Endoplasmic reticulum-resident selenoproteins as regulators of calcium signaling and homeostasis. Cell Calcium. 2018;70:76–86. doi: 10.1016/j.ceca.2017.05.001. PubMed PMID: WOS:000427523200009.

31. Anouar Y, Lihrmann I, Falluel-Morel A, Boukhzar L. Selenoprotein T is a key player in ER proteostasis, endocrine homeostasis and neuroprotection. Free Radical Biology and Medicine. 2018;127:145–52. doi: 10.1016/j.freeradbiomed.2018.05.076. PubMed PMID: WOS:000445890700015.

32. Tiengwe C, Brown A, Bangs JD. Unfolded Protein Response Pathways in Bloodstream-Form Trypanosoma brucei? Eukaryotic Cell. 2015;14(11):1094–101. doi: 10.1128/ec.00118.15. PubMed PMID: WOS:000363814000006.

33. Michaeli S. The response of trypanosomes and other eukaryotes to ER stress and the spliced leader RNA silencing (SLS) pathway in Trypanosoma brucei. Critical Reviews in Biochemistry and Molecular Biology. 2015;50(3):256–67. doi: 10.3109/10409238.2015.1042541. PubMed PMID: WOS:000361316400007.

34. Michaeli S. Spliced leader RNA silencing (SLS) - a programmed cell death pathway in Trypanosoma brucei that is induced upon ER stress. Parasites & Vectors. 2012;5. doi: 10.1186/1756-3305-5-107. PubMed PMID: WOS:000307339100002.

35. Bindereif A, Preusser C. ER stress: how trypanosomes deal with it. Trends in Parasitology. 2014;30(12):549–50. doi: 10.1016/j.pt.2014.10.006. PubMed PMID: WOS:000347140400001.

36. Faim LM, Rosa e Silva I, Bertacine Dias MV, Pereira HDM, Brandao-Neto J, Alves da Silva MT, et al. Crystallization and preliminary X-ray diffraction analysis of selenophosphate synthetases from Trypanosoma brucei and Leishmania major. Acta Crystallographica Section F-Structural Biology and Crystallization Communications. 2013;69:864–7. doi: 10.1107/s1744309113014632. PubMed PMID: WOS:000322512300009.

37. Li CL, Kappock TJ, Stubbe J, Weaver TM, Ealick SE. X-ray crystal structure of aminoimidazole ribonucleotide synthetase (PurM), from the Escherichia coli purine biosynthetic pathway at 2.5 angstrom resolution. Structure with Folding & Design. 1999;7(9):1155–66. doi: 10.1016/s0969-2126(99)80182-8. PubMed PMID: WOS:000082691000017.

38. Itoh Y, Sekine SI, Matsumoto E, Akasaka R, Takemoto C, Shirouzu M, et al. Structure of Selenophosphate Synthetase Essential for Selenium Incorporation into Proteins and RNAs. Journal of Molecular Biology. 2009;385(5):1456–69. doi: 10.1016/j.jmb.2008.08.042. PubMed PMID: WOS:000264383400011.

39. Noinaj N, Wattanasak R, Lee D-Y, Wally JL, Piszczek G, Chock PB, et al. Structural Insights into the Catalytic Mechanism of Escherichia coli Selenophosphate Synthetase. Journal of Bacteriology. 2012;194(2):499–508. doi: 10.1128/jb.06012-11. PubMed PMID: WOS:000298677400031.

40. Krissinel E, Henrick K. Inference of macromolecular assemblies from crystalline state. Journal of Molecular Biology. 2007;372(3):774–97. doi: 10.1016/j.jmb.2007.05.022. PubMed PMID: WOS:000249494800018.

41. Silva IR, Serrao VHB, Manzine LR, Faim LM, da Silva MTA, Makki R, et al. Formation of a Ternary Complex for Selenocysteine Biosynthesis in Bacteria. Journal of Biological Chemistry. 2015;290(49):29178–U103. doi: 10.1074/jbc.M114.613406. PubMed PMID: WOS:000366613400006.

42. Matsumoto E, Sekine S-i, Akasaka R, Otta Y, Katsura K, Inoue M, et al. Structure of an N-terminally truncated selenophosphate synthetase from Aquifex aeolicus. Acta Crystallographica Section F-Structural Biology and Crystallization Communications. 2008;64:453–8. doi: 10.1107/s1744309108012074. PubMed PMID: WOS:000256295000003.

43. Tobe R, Mihara H, Kurihara T, Esaki N. Identification of Proteins Interacting with Selenocysteine Lyase. Bioscience Biotechnology and Biochemistry. 2009;73(5):1230–2. doi: 10.1271/bbb.90065. PubMed PMID: WOS:000266620100049.

44. Small-Howard A, Morozova N, Stoytcheva Z, Forry EP, Mansell JB, Harney JW, et al. Supramolecular complexes mediate selenocysteine incorporation in vivo. Molecular and Cellular Biology. 2006;26(6):2337–46. doi: 10.1128/mcb.26.6.2337-2346.2006. PubMed PMID: WOS:000235915400028.

45. Goldshmidt H, Matas D, Kabi A, Carmi S, Hope R, Michaeli S. Persistent ER Stress Induces the Spliced Leader RNA Silencing Pathway (SLS), Leading to Programmed Cell Death in Trypanosoma brucei. Plos Pathogens. 2010;6(1). doi: 10.1371/journal.ppat.1000731. PubMed PMID: WOS:000274227100022.

46. Koumandou VL, Natesan SKA, Sergeenko T, Field MC. The trypanosome transcriptome is remodelled during differentiation but displays limited responsiveness within life stages. Bmc Genomics. 2008;9. doi: 10.1186/1471-2164-9-298. PubMed PMID: WOS:000257741900001.

47. Zeeshan HMA, Lee GH, Kim HR, Chae HJ. Endoplasmic Reticulum Stress and Associated ROS. International Journal of Molecular Sciences. 2016;17(3). doi: 10.3390/ijms17030327. PubMed PMID: WOS:000373712800087.

48. Omi R, Kurokawa S, Mihara H, Hayashi H, Goto M, Miyahara I, et al. Reaction Mechanism and Molecular Basis for Selenium/Sulfur Discrimination of Selenocysteine Lyase. Journal of Biological Chemistry. 2010;285(16):12133–9. doi: 10.1074/jbc.M109.084475. PubMed PMID: WOS:000276787100043.

49. Lacourciere GM, Mihara H, Kurihara T, Esaki N, Stadtman TC. Escherichia coli NifS-like proteins provide selenium in the pathway for the biosynthesis of selenophosphate. Journal of Biological Chemistry. 2000;275(31):23769–73. doi: 10.1074/jbc.M000926200. PubMed PMID: WOS:000088564200051.

50. Oudouhou F, Casu B, Puemi ASD, Sygusch J, Baron C. Analysis of Novel Interactions between Components of the Selenocysteine Biosynthesis Pathway, SEPHS1, SEPHS2, SEPSECS, and SECp43. Biochemistry. 2017;56(17):2261–70. doi: 10.1021/acs.biochem.6b01116. PubMed PMID: WOS:000400723300003.

51. Sherrer RL, Araiso Y, Aldag C, Ishitani R, Ho JML, Soell D, et al. C-terminal domain of archaeal O-phosphoseryl-tRNA kinase displays large-scale motion to bind the 7-bp D-stem of archaeal tRNA(Sec). Nucleic Acids Research. 2011;39(3):1034–41. doi: 10.1093/nar/gkq845. PubMed PMID: WOS:000287257500029.

52. Araiso Y, Palioura S, Ishitani R, Sherrer RL, O’Donoghue P, Yuan J, et al. Structural insights into RNA-dependent eukaryal and archaeal selenocysteine formation. Nucleic Acids Research. 2008;36(4):1187–99. doi: 10.1093/nar/gkm1122. PubMed PMID: WOS:000253857400019.

53. Ganichkin OM, Xu XM, Carlson BA, Mix H, Hatfield DL, Gladyshev VN, et al. Structure and catalytic mechanism of eukaryotic selenocysteine synthase. Journal of Biological Chemistry. 2008;283(9):5849–65. doi: 10.1074/jbc.M709342200. PubMed PMID: WOS:000253426700064.

54. Itoh Y, Broecker MJ, Sekine S-i, Hammond G, Suetsugu S, Soell D, et al. Decameric SelA.tRNA(Sec) Ring Structure Reveals Mechanism of Bacterial Selenocysteine Formation. Science. 2013;340(6128):75–8. doi: 10.1126/science.1229521. PubMed PMID: WOS:000317061100048.

55. Palioura S, Sherrer RL, Steitz TA, Soell D, Simonovic M. The Human SepSecS-tRNA(Sec) Complex Reveals the Mechanism of Selenocysteine Formation. Science. 2009;325(5938):321–5. doi: 10.1126/science.1173755. PubMed PMID: WOS:000268036600046.

56. Ridgley EL, Xiong ZH, Ruben L. Reactive oxygen species activate a Ca2+-dependent cell death pathway in the unicellular organism Trypanosoma brucei brucei. Biochemical Journal. 1999;340:33–40. doi: 10.1042/0264-6021:3400033. PubMed PMID: WOS:000080505300005.

57. Welburn SC, Macleod E, Figarella K, Duzensko M. Programmed cell death in African trypanosomes. Parasitology. 2006;132:S7–S18. doi: 10.1017/s0031182006000825. PubMed PMID: WOS:000242346400003.

58. Feige MJ, Hendershot LM. Disulfide bonds in ER protein folding and homeostasis. Current Opinion in Cell Biology. 2011;23(2):167–75. doi: 10.1016/j.ceb.2010.10.012. PubMed PMID: WOS:000290505000007.

59. Cabibbo A, Pagani M, Fabbri M, Rocchi M, Farmery MR, Bulleid NJ, et al. ERO1-L, a human protein that favors disulfide bond formation in the endoplasmic reticulum. Journal of Biological Chemistry. 2000;275(7):4827–33. doi: 10.1074/jbc.275.7.4827. PubMed PMID: WOS:000085378200043.

60. Pagani M, Fabbri M, Benedetti C, Fassio A, Pilati S, Bulleid NJ, et al. Endoplasmic reticulum oxidoreductin 1-L beta (ERO1-L beta), a human gene induced in the course of the unfolded protein response. Journal of Biological Chemistry. 2000;275(31):23685–92. doi: 10.1074/jbc.M003061200. PubMed PMID: WOS:000088564200040.

61. Dolai S, Adak S. Endoplasmic reticulum stress responses in Leishmania. Molecular and Biochemical Parasitology. 2014;197(1-2):1–8. doi: 10.1016/j.molbiopara.2014.09.002. PubMed PMID: WOS:000347129000001.

62. Strickler JE, Patton CL. TRYPANOSOMA-BRUCEI - EFFECT OF INHIBITION OF N-LINKED GLYCOSYLATION ON THE NEAREST NEIGHBOR ANALYSIS OF THE MAJOR VARIABLE SURFACE-COAT GLYCOPROTEIN. Molecular and Biochemical Parasitology. 1982;5(2):117–31. doi: 10.1016/0166-6851(82)90046-9. PubMed PMID: WOS:A1982NJ74000005.

63. Huang GZ, Vercesi AE, Docampo R. Essential regulation of cell bioenergetics in Trypanosoma brucei by the mitochondrial calcium uniporter. Nature Communications. 2013;4. doi: 10.1038/ncomms3865. PubMed PMID: WOS:000329395800001.

64. Addinsall AB, Wright CR, Andrikopoulos S, van der Poel C, Stupka N. Emerging roles of endoplasmic reticulum-resident selenoproteins in the regulation of cellular stress responses and the implications for metabolic disease. Biochemical Journal. 2018;475:1037–57. doi: 10.1042/bcj20170920. PubMed PMID: WOS:000431967100001.

65. Boukhzar L, Hamieh A, Cartier D, Tanguy Y, Alsharif I, Castex M, et al. Selenoprotein T Exerts an Essential Oxidoreductase Activity That Protects Dopaminergic Neurons in Mouse Models of Parkinson’s Disease. Antioxidants & Redox Signaling. 2016;24(11):557–74. doi: 10.1089/ars.2015.6478. PubMed PMID: WOS:000374645500001.

66. Cassago A, Rodrigues EM, Prieto EL, Gaston KW, Alfonzo JD, Iribar MP, et al. Identification of Leishmania selenoproteins and SECIS element. Molecular and Biochemical Parasitology. 2006;149(2):128–34. doi: 10.1016/j.molbiopara.2006.05.002. PubMed PMID: WOS:000240776600002.

67. Schwede A, Kramer S, Carrington M. How do trypanosomes change gene expression in response to the environment? Protoplasma. 2012;249(2):223–38. doi: 10.1007/s00709-011-0282-5. PubMed PMID: WOS:000305687700002.

68. Dolai S, Pal S, Yadav RK, Adak S. Endoplasmic Reticulum Stress-induced Apoptosis in Leishmania through Ca2+-dependent and Caspase-independent Mechanism. Journal of Biological Chemistry. 2011;286(15):13638–46. doi: 10.1074/jbc.M110.201889. PubMed PMID: WOS:000289282200082.

69. Izquierdo L, Atrih A, Rodrigues JA, Jones DC, Ferguson MAJ. Trypanosoma brucei UDP-Glucose:Glycoprotein Glucosyltransferase Has Unusual Substrate Specificity and Protects the Parasite from Stress. Eukaryotic Cell. 2009;8(2):230–40. doi: 10.1128/ec.00361-08. PubMed PMID: WOS:000263027200010.

70. Abhishek K, Das S, Kumar A, Kumar A, Kumar V, Saini S, et al. L-donovani induced Unfolded Protein Response delays host cell apoptosis in PERK dependent manner. Plos Neglected Tropical Diseases. 2018;12(7). doi: 10.1371/journal.pntd.0006646. PubMed PMID: WOS:000440495700046.

71. Walter P, Ron D. The Unfolded Protein Response: From Stress Pathway to Homeostatic Regulation. Science. 2011;334(6059):1081–6. doi: 10.1126/science.1209038. PubMed PMID: WOS:000297313900038.

72. Ashkenazy H, Abadi S, Martz E, Chay O, Mayrose I, Pupko T, et al. ConSurf 2016: an improved methodology to estimate and visualize evolutionary conservation in macromolecules. Nucleic Acids Research. 2016;44(W1):W344–W50. doi: 10.1093/nar/gkw408. PubMed PMID: WOS:000379786800057.

73. Schuck P. Size-distribution analysis of macromolecules by sedimentation velocity ultracentrifugation and Lamm equation modeling. Biophysical Journal. 2000;78(3):1606–19. PubMed PMID: WOS:000085697800047.

74. Geer LY, Marchler-Bauer A, Geer RC, Han LY, He J, He SQ, et al. The NCBI BioSystems database. Nucleic Acids Research. 2010;38:D492–D6. doi: 10.1093/nar/gkp858. PubMed PMID: WOS:000276399100079.

75. Sievers F, Wilm A, Dineen D, Gibson TJ, Karplus K, Li W, et al. Fast, scalable generation of high-quality protein multiple sequence alignments using Clustal Omega. Molecular Systems Biology. 2011;7. doi: 10.1038/msb.2011.75. PubMed PMID: WOS:000296652600007.

76. McCoy AJ, Grosse-Kunstleve RW, Adams PD, Winn MD, Storoni LC, Read RJ. Phaser crystallographic software. Journal of Applied Crystallography. 2007;40:658–74. doi: 10.1107/s0021889807021206. PubMed PMID: WOS:000248077500003.

77. Afonine PV, Grosse-Kunstleve RW, Echols N, Headd JJ, Moriarty NW, Mustyakimov M, et al. Towards automated crystallographic structure refinement with phenix.refine. Acta Crystallographica Section D-Biological Crystallography. 2012;68:352–67. doi: 10.1107/s0907444912001308. PubMed PMID: WOS:000302138400005.

78. Emsley P, Lohkamp B, Scott WG, Cowtan K. Features and development of Coot. Acta Crystallographica Section D-Biological Crystallography. 2010;66:486–501. doi: 10.1107/s0907444910007493. PubMed PMID: WOS:000275941300018.

79. Halliday C, Billington K, Wang Z, Madden R, Dean S, Sunter JD, Wheeler RJ. Cellular landmarks of *Trypanosoma brucei* and *Leishmania mexicana*. Molecular and Biochemical Parasitology 2019; 230, 24–36. doi:10.1016/j.molbiopara.2018.12.003.

80. Oberholzer M, Lopez MA, Ralston KS, Hill KL. Approaches for Functional Analysis of Flagellar Proteins in African Trypanosomes. Methods in Cell Biology. 2009; 21–55. Doi 10.1016/S0091-679X(08)93002-8.

81. Burkard GS, Jutzi P, Roditi I. Genome-wide RNAi screens in bloodstream form trypanosomes identify drug transporters. Molecular and Biochemical Parasitology. 2011; 175, 91–94. doi 10.1016/j.molbiopara.2010.09.002

## References

1. Kaufer A, Ellis J, Stark D, Barratt J (2017) The evolution of trypanosomatid taxonomy. Parasites & Vectors 10.

2. Franco JR, Simarro PP, Diarra A, Jannin JG (2014) Epidemiology of human African trypanosomiasis. Clinical epidemiology 6: 257–275.

3. Messenger LA, Miles MA, Bern C (2015) Between a bug and a hard place: Trypanosoma cruzi genetic diversity and the clinical outcomes of Chagas disease. Expert review of anti-infective therapy 13: 995–1029.

4. Kevric I, Cappel MA, Keeling JH (2015) New World and Old World Leishmania Infections: A Practical Review. Dermatologic clinics 33: 579–593.

5. Daniels J-P, Gull K, Wickstead B (2010) Cell Biology of the Trypanosome Genome. Microbiology and Molecular Biology Reviews 74: 552–569.

6. Guenzl A (2010) The Pre-mRNA Splicing Machinery of Trypanosomes: Complex or Simplified? Eukaryotic Cell 9.

7. Fairlamb AH, Cerami A (1992) METABOLISM AND FUNCTIONS OF TRYPANOTHIONE IN THE KINETOPLASTIDA. Annual Review of Microbiology 46: 695–729.

8. Krauth-Siegel RL, Comini MA (2008) Redox control in trypanosomatids, parasitic protozoa with trypanothione-based thiol metabolism. Biochimica Et Biophysica Acta-General Subjects 1780: 1236–1248.

9. Manta B, Comini M, Medeiros A, Hugo M, Trujillo M, et al. (2013) Trypanothione: A unique bis-glutathionyl derivative in trypanosomatids. Biochimica Et Biophysica Acta-General Subjects 1830: 3199–3216.

10. Steinbrenner H, Speckmann B, Klotz LO (2016) Selenoproteins: Antioxidant selenoenzymes and beyond. Archives of Biochemistry and Biophysics 595: 113–119.

11. Aeby E, Palioura S, Pusnik M, Marazzi J, Lieberman A, et al. (2009) The canonical pathway for selenocysteine insertion is dispensable in Trypanosomes. Proceedings of the National Academy of Sciences of the United States of America 106: 5088–5092.

12. Lobanov AV, Gromer S, Salinas G, Gladyshev VN (2006) Selenium metabolism in Trypanosoma: characterization of selenoproteomes and identification of a Kinetoplastida-specific selenoprotein. Nucleic Acids Research 34: 4012–4024.

13. Manhas R, Gowri VS, Madhubala R (2016) Leishmania donovani Encodes a Functional Selenocysteinyl-tRNA Synthase. Journal of Biological Chemistry 291: 1203–1220.

14. Serrao VHB, Silva IR, da Silva MTA, Scortecci JF, Fernandes AD, et al. (2018) The unique tRNA(Sec) and its role in selenocysteine biosynthesis. Amino Acids 50: 1145–1167.

15. Xu X-M, Carlson BA, Mix H, Zhang Y, Saira K, et al. (2007) Biosynthesis of selenocysteine on its tRNA in eukaryotes. Plos Biology 5: 96–105.

16. Bulteau AL, Chavatte L (2015) Update on Selenoprotein Biosynthesis. Antioxidants & Redox Signaling 23: 775–794.

17. da Silva MTA, Silva-Jardim I, Thiemann OH (2014) Biological Implications of Selenium and its Role in Trypanosomiasis Treatment. Current Medicinal Chemistry 21: 1772–1780.

18. Allmang C, Wurth L, Krol A (2009) The selenium to selenoprotein pathway in eukaryotes: More molecular partners than anticipated. Biochimica Et Biophysica Acta-General Subjects 1790: 1415–1423.

19. Collins R, Johansson A-L, Karlberg T, Markova N, van den Berg S, et al. (2012) Biochemical Discrimination between Selenium and Sulfur 1: A Single Residue Provides Selenium Specificity to Human Selenocysteine Lyase. Plos One 7.

20. Tobe R, Mihara H (2018) Delivery of selenium to selenophosphate synthetase for selenoprotein biosynthesis. Biochimica Et Biophysica Acta-General Subjects 1862: 2433–2440.

21. Poliak P, Van Hoewyk D, Obornik M, Zikova A, Stuart KD, et al. (2010) Functions and cellular localization of cysteine desulfurase and selenocysteine lyase in Trypanosoma brucei. Febs Journal 277: 383–393.

22. Wang KT, Wang J, Li LF, Su XD (2009) Crystal Structures of Catalytic Intermediates of Human Selenophosphate Synthetase 1. Journal of Molecular Biology 390: 747–759.

23. Lobanov AV, Hatfield DL, Gladyshev VN (2008) Selenoproteinless animals: Selenophosphate synthetase SPS1 functions in a pathway unrelated to selenocysteine biosynthesis. Protein Science 17: 176–182.

24. Xu XM, Carlson BA, Irons R, Mix H, Zhong NX, et al. (2007) Selenophosphate synthetase 2 is essential for selenoprotein biosynthesis. Biochemical Journal 404: 115–120.

25. Na J, Jung J, Bang J, Lu Q, Carlson BA, et al. (2018) Selenophosphate synthetase 1 and its role in redox homeostasis, defense and proliferation. Free Radical Biology and Medicine 127: 190–197.

26. Sculaccio SA, Rodrigues EM, Cordeiro AT, Magalhaes A, Braga AL, et al. (2008) Selenocysteine incorporation in Kinetoplastid: Selenophosphate synthetase (SELD) from Leishmania major and Trypanosoma brucei. Molecular and Biochemical Parasitology 162: 165–171.

27. Aeby E, Seidel V, Schneider A (2009) The selenoproteome is dispensable in bloodstream forms of Trypanosoma brucei. Molecular and Biochemical Parasitology 168: 191–193.

28. Bonilla M, Krull E, Irigoin F, Salinas G, Comini MA (2016) Selenoproteins of African trypanosomes are dispensable for parasite survival in a mammalian host. Molecular and Biochemical Parasitology 206: 13–19.

29. Costa FC, Oliva MAV, de Jesus TCL, Schenkman S, Thiemann OH (2011) Oxidative stress protection of Trypanosomes requires selenophosphate synthase. Molecular and Biochemical Parasitology 180: 47–50.

30. Pitts MW, Hoffmann PR (2018) Endoplasmic reticulum-resident selenoproteins as regulators of calcium signaling and homeostasis. Cell Calcium 70: 76–86.

31. Anouar Y, Lihrmann I, Falluel-Morel A, Boukhzar L (2018) Selenoprotein T is a key player in ER proteostasis, endocrine homeostasis and neuroprotection. Free Radical Biology and Medicine 127: 145–152.

32. Tiengwe C, Brown A, Bangs JD (2015) Unfolded Protein Response Pathways in Bloodstream-Form Trypanosoma brucei? Eukaryotic Cell 14: 1094–1101.

33. Michaeli S (2015) The response of trypanosomes and other eukaryotes to ER stress and the spliced leader RNA silencing (SLS) pathway in Trypanosoma brucei. Critical Reviews in Biochemistry and Molecular Biology 50: 256–267.

34. Michaeli S (2012) Spliced leader RNA silencing (SLS) - a programmed cell death pathway in Trypanosoma brucei that is induced upon ER stress. Parasites & Vectors 5.

35. Bindereif A, Preusser C (2014) ER stress: how trypanosomes deal with it. Trends in Parasitology 30: 549–550.

36. Faim LM, Rosa e Silva I, Bertacine Dias MV, Pereira HDM, Brandao-Neto J, et al. (2013) Crystallization and preliminary X-ray diffraction analysis of selenophosphate synthetases from Trypanosoma brucei and Leishmania major. Acta Crystallographica Section F-Structural Biology and Crystallization Communications 69: 864–867.

37. Li CL, Kappock TJ, Stubbe J, Weaver TM, Ealick SE (1999) X-ray crystal structure of aminoimidazole ribonucleotide synthetase (PurM), from the Escherichia coli purine biosynthetic pathway at 2.5 angstrom resolution. Structure with Folding & Design 7: 1155–1166.

38. Itoh Y, Sekine SI, Matsumoto E, Akasaka R, Takemoto C, et al. (2009) Structure of Selenophosphate Synthetase Essential for Selenium Incorporation into Proteins and RNAs. Journal of Molecular Biology 385: 1456–1469.

39. Noinaj N, Wattanasak R, Lee D-Y, Wally JL, Piszczek G, et al. (2012) Structural Insights into the Catalytic Mechanism of Escherichia coli Selenophosphate Synthetase. Journal of Bacteriology 194: 499–508.

40. Krissinel E, Henrick K (2007) Inference of macromolecular assemblies from crystalline state. Journal of Molecular Biology 372: 774–797.

41. Silva IR, Serrao VHB, Manzine LR, Faim LM, da Silva MTA, et al. (2015) Formation of a Ternary Complex for Selenocysteine Biosynthesis in Bacteria. Journal of Biological Chemistry 290: 29178–U29103.

42. Matsumoto E, Sekine S-i, Akasaka R, Otta Y, Katsura K, et al. (2008) Structure of an N-terminally truncated selenophosphate synthetase from Aquifex aeolicus. Acta Crystallographica Section F-Structural Biology and Crystallization Communications 64: 453–458.

43. Tobe R, Mihara H, Kurihara T, Esaki N (2009) Identification of Proteins Interacting with Selenocysteine Lyase. Bioscience Biotechnology and Biochemistry 73: 1230–1232.

44. Small-Howard A, Morozova N, Stoytcheva Z, Forry EP, Mansell JB, et al. (2006) Supramolecular complexes mediate selenocysteine incorporation in vivo. Molecular and Cellular Biology 26: 2337–2346.

45. Goldshmidt H, Matas D, Kabi A, Carmi S, Hope R, et al. (2010) Persistent ER Stress Induces the Spliced Leader RNA Silencing Pathway (SLS), Leading to Programmed Cell Death in Trypanosoma brucei. Plos Pathogens 6.

46. Koumandou VL, Natesan SKA, Sergeenko T, Field MC (2008) The trypanosome transcriptome is remodelled during differentiation but displays limited responsiveness within life stages. Bmc Genomics 9.

47. Zeeshan HMA, Lee GH, Kim HR, Chae HJ (2016) Endoplasmic Reticulum Stress and Associated ROS. International Journal of Molecular Sciences 17.

48. Omi R, Kurokawa S, Mihara H, Hayashi H, Goto M, et al. (2010) Reaction Mechanism and Molecular Basis for Selenium/Sulfur Discrimination of Selenocysteine Lyase. Journal of Biological Chemistry 285: 12133–12139.

49. Lacourciere GM, Mihara H, Kurihara T, Esaki N, Stadtman TC (2000) Escherichia coli NifS-like proteins provide selenium in the pathway for the biosynthesis of selenophosphate. Journal of Biological Chemistry 275: 23769–23773.

50. Oudouhou F, Casu B, Puemi ASD, Sygusch J, Baron C (2017) Analysis of Novel Interactions between Components of the Selenocysteine Biosynthesis Pathway, SEPHS1, SEPHS2, SEPSECS, and SECp43. Biochemistry 56: 2261–2270.

51. Sherrer RL, Araiso Y, Aldag C, Ishitani R, Ho JML, et al. (2011) C-terminal domain of archaeal O-phosphoseryl-tRNA kinase displays large-scale motion to bind the 7-bp D-stem of archaeal tRNA(Sec). Nucleic Acids Research 39: 1034–1041.

52. Araiso Y, Palioura S, Ishitani R, Sherrer RL, O’Donoghue P, et al. (2008) Structural insights into RNA-dependent eukaryal and archaeal selenocysteine formation. Nucleic Acids Research 36: 1187–1199.

53. Ganichkin OM, Xu XM, Carlson BA, Mix H, Hatfield DL, et al. (2008) Structure and catalytic mechanism of eukaryotic selenocysteine synthase. Journal of Biological Chemistry 283: 5849–5865.

54. Itoh Y, Broecker MJ, Sekine S-i, Hammond G, Suetsugu S, et al. (2013) Decameric SelA.tRNA(Sec) Ring Structure Reveals Mechanism of Bacterial Selenocysteine Formation. Science 340: 75–78.

55. Palioura S, Sherrer RL, Steitz TA, Soell D, Simonovic M (2009) The Human SepSecS-tRNA(Sec) Complex Reveals the Mechanism of Selenocysteine Formation. Science 325: 321–325.

56. Ridgley EL, Xiong ZH, Ruben L (1999) Reactive oxygen species activate a Ca2+-dependent cell death pathway in the unicellular organism Trypanosoma brucei brucei. Biochemical Journal 340: 33–40.

57. Welburn SC, Macleod E, Figarella K, Duzensko M (2006) Programmed cell death in African trypanosomes. Parasitology 132: S7–S18.

58. Feige MJ, Hendershot LM (2011) Disulfide bonds in ER protein folding and homeostasis. Current Opinion in Cell Biology 23: 167–175.

59. Cabibbo A, Pagani M, Fabbri M, Rocchi M, Farmery MR, et al. (2000) ERO1-L, a human protein that favors disulfide bond formation in the endoplasmic reticulum. Journal of Biological Chemistry 275: 4827–4833.

60. Pagani M, Fabbri M, Benedetti C, Fassio A, Pilati S, et al. (2000) Endoplasmic reticulum oxidoreductin 1-L beta (ERO1-L beta), a human gene induced in the course of the unfolded protein response. Journal of Biological Chemistry 275: 23685–23692.

61. Dolai S, Adak S (2014) Endoplasmic reticulum stress responses in Leishmania. Molecular and Biochemical Parasitology 197: 1–8.

62. Strickler JE, Patton CL (1982) TRYPANOSOMA-BRUCEI - EFFECT OF INHIBITION OF N-LINKED GLYCOSYLATION ON THE NEAREST NEIGHBOR ANALYSIS OF THE MAJOR VARIABLE SURFACE-COAT GLYCOPROTEIN. Molecular and Biochemical Parasitology 5: 117–131.

63. Huang GZ, Vercesi AE, Docampo R (2013) Essential regulation of cell bioenergetics in Trypanosoma brucei by the mitochondrial calcium uniporter. Nature Communications 4.

64. Addinsall AB, Wright CR, Andrikopoulos S, van der Poel C, Stupka N (2018) Emerging roles of endoplasmic reticulum-resident selenoproteins in the regulation of cellular stress responses and the implications for metabolic disease. Biochemical Journal 475: 1037–1057.

65. Boukhzar L, Hamieh A, Cartier D, Tanguy Y, Alsharif I, et al. (2016) Selenoprotein T Exerts an Essential Oxidoreductase Activity That Protects Dopaminergic Neurons in Mouse Models of Parkinson’s Disease. Antioxidants & Redox Signaling 24: 557–574.

66. Cassago A, Rodrigues EM, Prieto EL, Gaston KW, Alfonzo JD, et al. (2006) Identification of Leishmania selenoproteins and SECIS element. Molecular and Biochemical Parasitology 149: 128–134.

67. Schwede A, Kramer S, Carrington M (2012) How do trypanosomes change gene expression in response to the environment? Protoplasma 249: 223–238.

68. Dolai S, Pal S, Yadav RK, Adak S (2011) Endoplasmic Reticulum Stress-induced Apoptosis in Leishmania through Ca2+-dependent and Caspase-independent Mechanism. Journal of Biological Chemistry 286: 13638–13646.

69. Izquierdo L, Atrih A, Rodrigues JA, Jones DC, Ferguson MAJ (2009) Trypanosoma brucei UDP-Glucose:Glycoprotein Glucosyltransferase Has Unusual Substrate Specificity and Protects the Parasite from Stress. Eukaryotic Cell 8: 230–240.

70. Abhishek K, Das S, Kumar A, Kumar A, Kumar V, et al. (2018) L-donovani induced Unfolded Protein Response delays host cell apoptosis in PERK dependent manner. Plos Neglected Tropical Diseases 12.

71. Walter P, Ron D (2011) The Unfolded Protein Response: From Stress Pathway to Homeostatic Regulation. Science 334: 1081–1086.

72. Ashkenazy H, Abadi S, Martz E, Chay O, Mayrose I, et al. (2016) ConSurf 2016: an improved methodology to estimate and visualize evolutionary conservation in macromolecules. Nucleic Acids Research 44: W344–W350.

73. Schuck P (2000) Size-distribution analysis of macromolecules by sedimentation velocity ultracentrifugation and Lamm equation modeling. Biophysical Journal 78: 1606–1619.

74. Geer LY, Marchler-Bauer A, Geer RC, Han LY, He J, et al. (2010) The NCBI BioSystems database. Nucleic Acids Research 38: D492–D496.

75. Sievers F, Wilm A, Dineen D, Gibson TJ, Karplus K, et al. (2011) Fast, scalable generation of high-quality protein multiple sequence alignments using Clustal Omega. Molecular Systems Biology 7.

76. McCoy AJ, Grosse-Kunstleve RW, Adams PD, Winn MD, Storoni LC, et al. (2007) Phaser crystallographic software. Journal of Applied Crystallography 40: 658–674.

77. Afonine PV, Grosse-Kunstleve RW, Echols N, Headd JJ, Moriarty NW, et al. (2012) Towards automated crystallographic structure refinement with phenix.refine. Acta Crystallographica Section D-Biological Crystallography 68: 352–367.

78. Emsley P, Lohkamp B, Scott WG, Cowtan K (2010) Features and development of Coot. Acta Crystallographica Section D-Biological Crystallography 66: 486–501.

79. Zhao HY, Piszczek G, Schuck P (2015) SEDPHAT - A platform for global ITC analysis and global multi-method analysis of molecular interactions. Methods 76: 137–148.

